# Chromosome-scale genome assemblies of duckweeds provide insights into genomic plasticity, aquatic adaptation and morphological reduction

**DOI:** 10.64898/2026.04.27.721116

**Authors:** Mümin Ibrahim Tek, Brian Boyle, Eric Normandeau, Laurie Lecomte, Mourad Sabri, Davoud Torkamaneh

**Affiliations:** Département de phytologie, Université Laval, Québec City, Quebec, Canada; Institut de Biologie Intégrative et des Systèmes (IBIS), Université Laval, Québec City, Quebec, Canada; Centre de recherche et d’innovation sur les végétaux (CRIV) Université Laval, Québec City, Quebec, Canada; Institute Intelligence and Data (IID), Université Laval, Québec City, Quebec, Canada; Institute of Nutrition and Functional Foods (INAF), Université Laval, Québec City, Quebec, Canada; Plateforme d’Analyses Génomiques de l’IBIS (Institut de Biologie Intégrative et des Systèmes), Université Laval, Québec, G1V 0A6, Canada; Plateforme de Bio-Informatique de l’IBIS (Institut de Biologie Intégrative et des Systèmes), Université Laval, Québec, G1V 0A6, Canada; Aplantex, Montréal, Québec, Canada

**Keywords:** duckweed, evolution, Lemnaceae, comparative genomics, allopolyploidy, terrestrial-to-aquatic transition, gene family contraction, effector-triggered immunity, secondary metabolite biosynthesis, convergent evolution

## Abstract

**Background:** Duckweeds (Lemnaceae) present a striking example of convergent genome evolution following the return from land to water. As the smallest and fastest-growing angiosperms, they exhibit extreme morphological reduction yet retain remarkable genomic plasticity through recurrent interspecific hybridisation, chromosomal rearrangements, and selective gene-family remodelling. The genomic mechanisms that distinguish this secondarily aquatic lifestyle from terrestrial ancestors, and whether these changes are convergent with other aquatic lineages such as seagrasses, have remained incompletely resolved.

**Results:** Here we report chromosome-scale genome assemblies for four duckweed species, *Spirodela polyrhiza*, *Lemna minuta*, *Lemna japonica*, and *Lemna aequinoctialis*, generated with PacBio HiFi long reads and Omni-C chromatin conformation capture. These assemblies include the first genomic characterisation of an unresolved hybrid lineage (*L. aequinoctialis* ×) that harbours a previously uncharacterised 3.5 Mb reciprocal translocation between subgenomes, as well as confirmation of the allodiploid origin of L. japonica. Comparative phylogenomics with land plants and the seagrass *Zostera marina* reveals a coherent, non-random programme of gene loss: effector-triggered immunity (ETI) components (EDS1 and PAD4) and the high-affinity nitrate transporter NRT2 are convergently absent across duckweeds and *Zostera marina*, consistent with relaxed pathogen pressure and abundant dissolved nutrients in aquatic habitats. In contrast, secondary-metabolite biosynthesis pathways for flavonoids, anthocyanins, flavones and flavonols are retained or expanded despite overall genome compaction.

**Conclusion:** These findings illustrate how the return to aquatic environments following terrestrialisation shaped duckweed genome evolution through convergent gene loss and selective pathway retention, and provide high-quality genomic resources to support future research in plant evolutionary biology and biotechnology.

## Background

Aquatic plants have a special position in the evolution of angiosperms owing to the transition from land back to water following the terrestrialisation of plants [1]. Duckweeds are distinguished by their simplified morphology and pervasive gene loss across diverse aquatic and terrestrial plant lineages [2]. They diverged from basal Alismatales lineages ∼120 million years ago (Mya) during the Cretaceous period and later from basal Araceae ∼110 Mya, followed by family diversification around 63 Mya [3,4]. Duckweeds are the *smallest and fastest-growing flowering plants* in the green lineage and are frequently considered invasive species in aquatic ecosystems [5,6].

Duckweeds comprise four main genera: *Spirodela*, *Lemna*, *Wolffia*, and *Wolffiella*, which display considerable morphological and genomic diversity [7]. Their genomes reflect a combination of polyploidy, recurrent interspecific hybridisation, and convergent reductive evolution. *S. polyrhiza* and *S. intermedia* possess the most compact genomes in duckweeds, with genome sizes of approximately 120–160 Mb [8]. In contrast, species of *Lemna*, such as *L. japonica* or *L. aequinoctialis*, have larger and more complex genomes ranging from approximately 200 to 950 Mb [3], whilst *Wolffia* species have genomes of around 430 Mb [9]. Intraspecific variation in ploidy and chromosome number is common; for example, *L. aequinoctialis* includes diploid (2n = 40), tetraploid (2n = 80), and triploid clones [8,10]. Interspecific hybridisation is also common in duckweeds. *L. japonica* is a hybrid between *L. minor* and *L. turionifera* [11], and *L. mediterranea* represents a hybrid derived from *L. minor* and *L. gibba* [12]. *L. aequinoctialis* can also hybridise with *L. perpusilla*, forming diploid or tetraploid hybrids known as *L. aoukikusa* [13]. Backcross hybrids between *L. aequinoctialis* and *L. aoukikusa* have also been reported [10].

Duckweeds also exhibit notable morphological variation associated with gene family contraction across different species [14]. *Spirodela* and *Landoltia* possess multiple roots with aerenchyma, whereas *Lemna* species typically have a single root lacking aerenchyma, in contrast, *Wolffia* and *Wolffiella* species are rootless [2]. Reductions in genes associated with phytohormone pathways, stomatal closure, and lignin biosynthesis have similarly been reported in *S. polyrhiza*, *L. minor*, and *L. punctata* [4].

Duckweeds have attracted considerable interest as platforms for plant biotechnology owing to their rapid clonal growth, small size, suitability for aquatic cultivation, and low production costs, making them particularly promising for molecular farming [15]. Various recombinant proteins have been successfully produced in duckweeds, including the M2 matrix protein from avian influenza virus (H5N1) in *L. minor* and hirudin in *W. arrhiza* [16,17]. Metabolic engineering has also demonstrated the potential of duckweeds for lipid production, with genetically modified *L. japonica* accumulating triacylglycerol (TAG) at levels comparable to soybean and oil palm [18]. Notably, a duckweed-based edible vaccine has been shown to confer complete protection against avian infectious bronchitis virus, representing a significant milestone in plant-based edible vaccine development for poultry [19]. As a result, duckweeds have emerged as an attractive model for both plant biology and biotechnological applications [20].

Despite growing interest in duckweeds as model systems for plant biology and biotechnology, genomic resources across Lemnaceae remain limited [21]. Generating high-quality genome assemblies across diverse lineages is therefore essential to better understand genome evolution and functional diversification within this family [22]. Here, we present high-quality *de novo* genome assemblies for four duckweed species: *S. polyrhiza*, characterised by a compact genome; *L. minuta*, a small-genome invasive *Lemna* species; *L. aequinoctialis* ×, representing a lineage with extensive genomic variation and hybridisation; and *L. japonica*, a hybrid with a larger and more complex genome. Notably, we identify *L. aequinoctialis* D3 for the first time as an unresolved hybrid lineage (hereafter referred to as ’*L. aequinoctialis ×*’ to denote its unresolved hybrid ancestry). In addition, our *L. minuta* assembly represents a substantial improvement over the previously published assembly [23] for this invasive species. These high-quality assemblies, combined with comprehensive annotations and comparative analyses provide new insights into genome evolution, structural diversity, and metabolic pathway composition across duckweeds.

## Results

### Chromosome-level genome assembly

We generated 1,987,182–4,030,490 PacBio HiFi reads (∼21.8–41.9 Gbp) for *S. polyrhiza*, *L. minuta*, *L. japonica*, and *L. aequinoctialis*, together with 316,277,047–399,958,802 Omni-C reads (∼47.4–59.9 Gbp). GenomeScope profiling [24] of HiFi and Omni-C reads (k = 21) indicated diploid genome organisation for all species (Figure S1). *S. polyrhiza* exhibited low heterozygosity (0.44–0.83%) and the lowest repetitive fraction (∼6.1%), indicative of a compact genome. In contrast, *L. minuta* showed higher heterozygosity (∼2.52%) and a repetitive fraction of ∼39.3–40%. *L. japonica* exhibited ∼35.8–37.7% repetitive content but very low heterozygosity (∼0.48%). *L. aequinoctialis* × had the highest repeat proportion (∼43.6–46.3%) and the highest heterozygosity (6.77–7.43%) among the species analysed, based on GenomeScope profiles of both HiFi and Omni-C datasets.

Genome assemblies were generated from PacBio HiFi reads, followed by decontamination and haplotig purging, and subsequent scaffolding with Omni-C data and manual curation. *L. minuta* exhibited the smallest genome among the *Lemna* species, with a total assembly size of 312.7 Mb and an N50 of 15.9 Mb (Table 1). Omni-C contact maps revealed a clear and continuous diagonal signal, supporting chromosome-scale assembly into 21 chromosomes without evidence of misjoins or misassemblies (Figure 1a). *S. polyrhiza* had the smallest genome among all assembled species (137 Mb, N50 = 7.78 Mb) and similarly displayed a clean diagonal pattern in the contact map, supporting 20 highly contiguous and well-resolved chromosomes (Figure 1b).

**Table 1.** Assessment of genome assembly quality and completeness using QUAST and BUSCO.

| Species | *L. aequinoctialis* × | *L. japonica* | *L. minuta* | *S. polyrhiza* |
| --- | --- | --- | --- | --- |
| **Total size (bp)** | 796,331,008 | 748,095,292 | 312,771,632 | 137,080,990 |
| **Scaffold number** | 61 | 75 | 94 | 62 |
| **Scaffold N50** | 20,421,085 | 18,919,663 | 15,923,464 | 7,788,018 |
| **Number of chromosomes** | 42 | 42 | 21 | 20 |
| **Proportion anchored into chromosomes** | 0.999 | 0.999 | 0.996 | 0.998 |
| **BUSCO\*** | S:5.41%, 23 | S:6.35%, 27 | S:98.78%, 409 | S:96.70%, 411 |
|  | D:93.64%, 398 | D:92.23%, 392 | D:1.88%, 8 | D:1.17%, 5 |
|  | F:0.47%, 2 | F:0.94%, 4 | F:1.41%, 6 | F:1.64%, 7 |
|  | M:0.47%, 2 | M:0.47%, 2 | M:0.47%, 2 | M:0.47%, 2 |
|  | N:425 | N:425 | N:425 | N:425 |

S: Single, D: Duplication, F: Fragmented, M: Missing, and N: Number of genes in database. \* High BUSCO duplication scores in *L. japonica* and *L. aequinoctialis* × are consistent with their hybrid origin and the presence of two near-identical subgenomes, rather than reflecting assembly artefacts.

**Figure 1.**
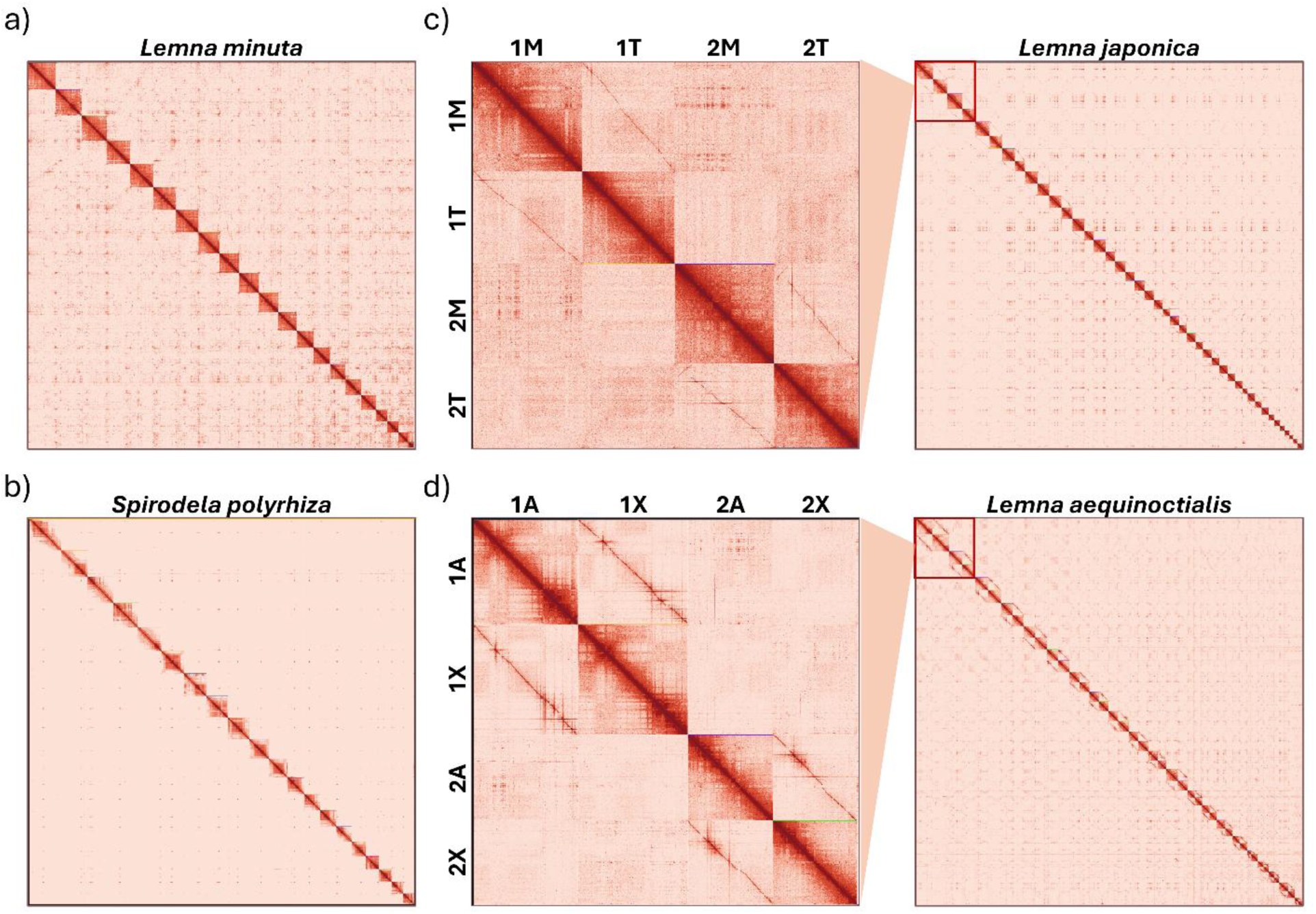
Omni-C contact maps illustrating chromatin organisation and genome structure in duckweed species following manual curation. Each pixel represents the normalised contact frequency between genomic loci, with colour intensity reflecting interaction strength. **a)** *L. minuta* displays a strong and continuous diagonal pattern, indicative of a well-resolved diploid genome organisation. **b)** *S. polyrhiza* similarly shows a clear diagonal signal, reflecting a compact and well-organised diploid genome with interactions largely confined within individual chromosomes**. c)** *L. japonica*, an allodiploid hybrid, exhibits off-diagonal signal between distinct and homeologous chromosomes, such as the first two homeologous pairs depicted in the zoomed-in region, in each of which one chromosome is derived from *L. minor* (chromosomes 1M, 2M), and the other from *L. turionifera* (1T, 2T). **d)** *L. aequinoctialis* × displays a similar contact map structure, with block-like off-diagonal interaction patterns indicating two subgenomes derived from *L. aequinoctialis* (1A, 2A) and an unknown parental lineage (1X, 2X), supporting a hybrid genome organisation.

Direct comparison with the most recent assemblies shows that our *L. minuta* assembly contains fewer scaffolds (94) than GCA_964341035 (997 scaffolds) and GCA_024174645.1 (2,381 scaffolds). The N50 of the new assembly is 15.92 Mbp, compared with 14.91 Mbp for GCA_964341035. In addition, the new assembly exhibits a 2–4% increase in total BUSCO completeness and a 7–10% increase in single-copy BUSCOs (File S1). Similarly, new *S. polyrhiza* assembly (N50 = 7.78 Mbp) shows a comparable N50 to GCA_024713555 (7.83 Mbp), while having substantially fewer scaffolds, with a 56.64% reduction in contig number. Moreover, the new assembly shows a 7% increase in single-copy BUSCOs.

In contrast, the assemblies of *L. aequinoctialis* × and *L. japonica* revealed more complex contact map patterns. The *L. aequinoctialis* × assembly spanned 796 Mb with an N50 of 20.4 Mb across 68 contigs, including 46 primary scaffolds larger than 10 Mb that accounted for 99.9% of the assembly, of which 42 were identified as putative chromosomes (Table 1). Omni-C contact maps of the *L. aequinoctialis* × D3 assembly revealed prominent off-diagonal interaction signals forming parallel blocks between chromosome pairs (Figure 1d), a pattern diagnostic of allodiploid hybrid genomes. A similar pattern was observed in *L. japonica* (748 Mb, N50 = 18.9 Mb) (Figure 1c). Both genomes exhibited a high proportion of duplicated BUSCO hits (93% and 92%, respectively), corroborating their hybrid origin (Table 1).

### Genome Structure and Synteny Among Duckweeds

Whole genome synteny analysis revealed clear subgenome relationships consistent with hybrid origins. In *L. japonica*, all chromosomes showed near-perfect one-to-one correspondence with chromosomes from parental assemblies of either *L. minor* (7210) and *L. turionifera* (9434) [3], respectively (Figure 2a). The total assembled size of *L. japonica* (746.8 Mb) closely matched the sum of its inferred parents (362.1 Mb + 405.9 Mb), confirming its allodiploid origin. Within the genus Spirodela, *S. polyrhiza* and *S. intermedia* exhibited extensive macrosynteny, with >95% of chromosomal regions aligned in conserved blocks. In contrast, synteny between *L. minuta* or *L. aequinoctialis* × and other Lemna species was markedly lower, reflecting greater structural divergence.

**Figure 2.**
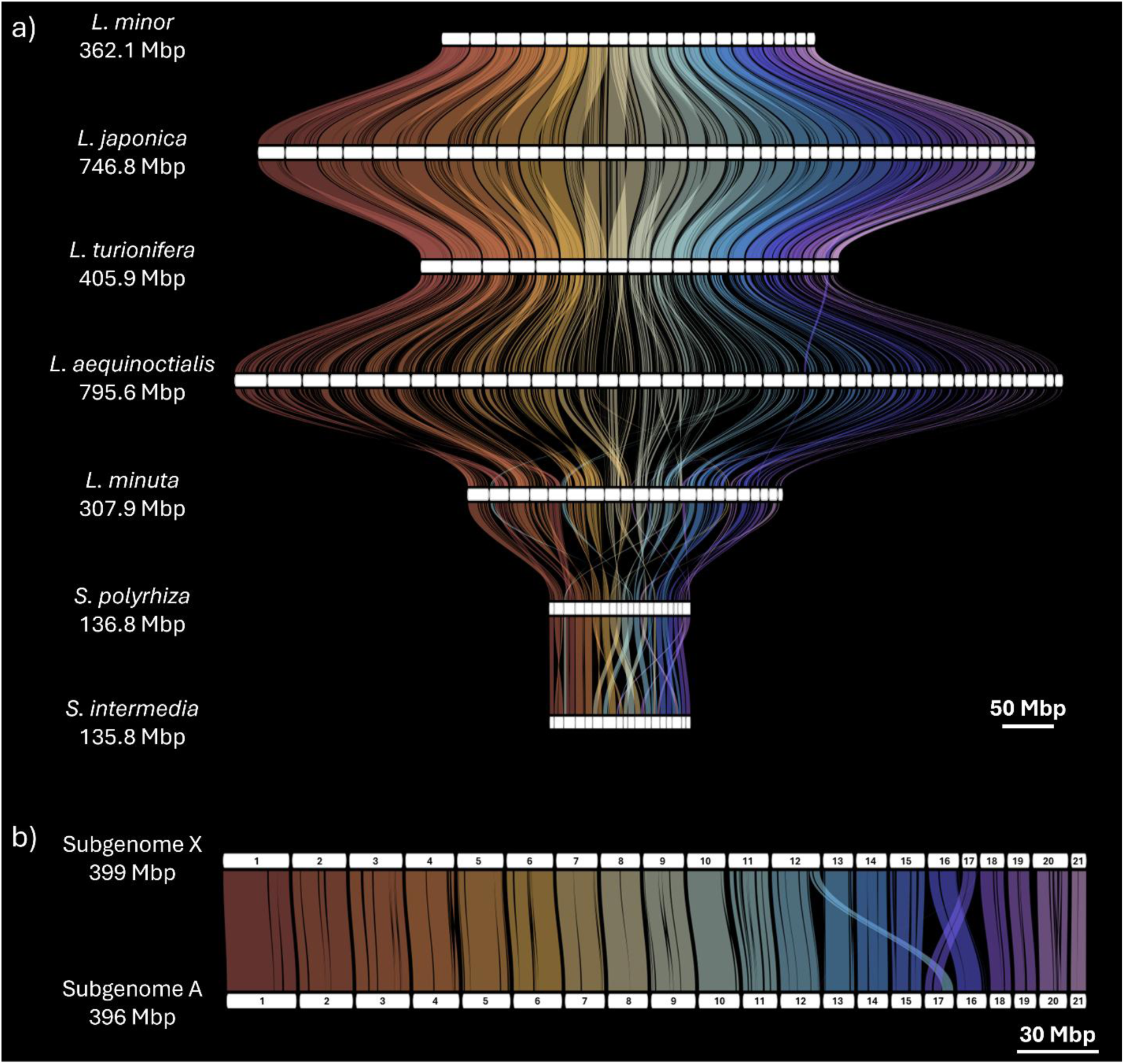
Genome-wide synteny across duckweed species and subgenome comparison of *Lemna aequinoctialis* × using DEEPSPACE. **a)** Whole-genome synteny comparison across seven duckweed genomes: *L. minor* 7210 (362.1 Mb), *L. japonica* (746.8 Mb), *L. turionifera* 9434 (405.9 Mb), *L. aequinoctialis* × (795.6 Mb), *L. minuta* (307.9 Mb), *S. polyrhiza* (136.8 Mb), and *S. intermedia* 8410 (135.8 Mb). White rectangles represent individual chromosomes, and coloured ribbons indicate syntenic relationships between genomes. **b)** Pairwise synteny between the two subgenomes of *L. aequinoctialis* ×, subgenome A (399 Mb) and subgenome X (396 Mb), revealing a high degree of collinearity between homologous chromosomes. A notable exception is observed between chromosomes 12 and 17, where a reciprocal translocation differentiates the two subgenomes, suggesting that this rearrangement occurred following hybridisation. Raw synteny maps for all species and for the *L. aequinoctialis* × subgenomes are provided in Files S6 and S7, respectively.

For *L. aequinoctialis* ×, intra-subgenome synteny identified a single large-scale rearrangement: a reciprocal translocation of 3.5 Mb between the telomeric region of chromosome 12X (subgenome X) and chromosome 17A (subgenome A) (Figure 2b). This rearrangement was independently confirmed by Omni-C contact maps showing collinearity between the translocated segments of chromosomes 12 and 17 (Figure S2). Supporting evidence for the unresolved hybrid ancestry came from rDNA analysis: the ITS1–5.8S–ITS2 region extracted from the assembled genome was identical to sequences from multiple known hybrid lineages, including *L. aequinoctialis* × *L. aoukikusa* strains 9668, NBP3, and NGY128, and *L. aequinoctialis* × *L. perpusilla* strain 7006 (File S8). We therefore designate this lineage *L. aequinoctialis* × to indicate its unresolved hybrid ancestry. No other major inter-subgenome rearrangements were detected.

### Genome Annotation

Gene prediction identified 17,588 protein-coding genes in *S. polyrhiza*, 22,481 in *L. minuta*, 55,470 in *L. japonica*, and 56,962 in *L. aequinoctialis* × genome assemblies. Comparative analysis revealed that *S. polyrhiza* possesses a compact genome with the highest gene density (42.63%), relatively low TE and repeats (20.91%) content, compared with *L. minuta*, which displayed the opposite pattern with a much lower proportion of coding sequence (19.39%) and higher TE and repeats (65.47%). Gene-rich regions tended to coincide with higher GC (≥42%) content, but gene density decreased in low-GC regions (≤40%), a pattern clearly observed in centromeric regions of *S. polyrhiza*. For instance, centromeric segments of chromosomes 7 (37.4% GC) and 8 (37.9% GC) corresponded to reduced gene density with 51.3 gene/1Mbp and 77 gene/1Mbp, while average was 142.92 gene/1Mbp within genome.

In contrast, the larger and hybrid genomes of *L. japonica* (19.60%) and *L. aequinoctialis* × (18.73%) showed gene densities comparable to *L. minuta* but were distinguished by substantially higher TE content with 72.5% and 73.2%, respectively. Beyond differences in total TE abundance, the relative composition of LTR retrotransposon families also varied between *S. polyrhiza* and *Lemna* species. In *S. polyrhiza*, LTR/Gypsy elements (6.02%) were more abundant than LTR/Copia elements (4.39%) among all annotated repeats (Figure 3). A similar Gypsy-dominant pattern was observed in *L. minuta*, where Gypsy elements (17.77%) substantially outnumbered Copia elements (9.73%). In contrast, this relationship was reversed in *L. aequinoctialis* × (Copia 19.29%, Gypsy 12.32%) and *L. japonica* (Copia 20.58%, Gypsy 18.76%) (File S2).

**Figure 3.**
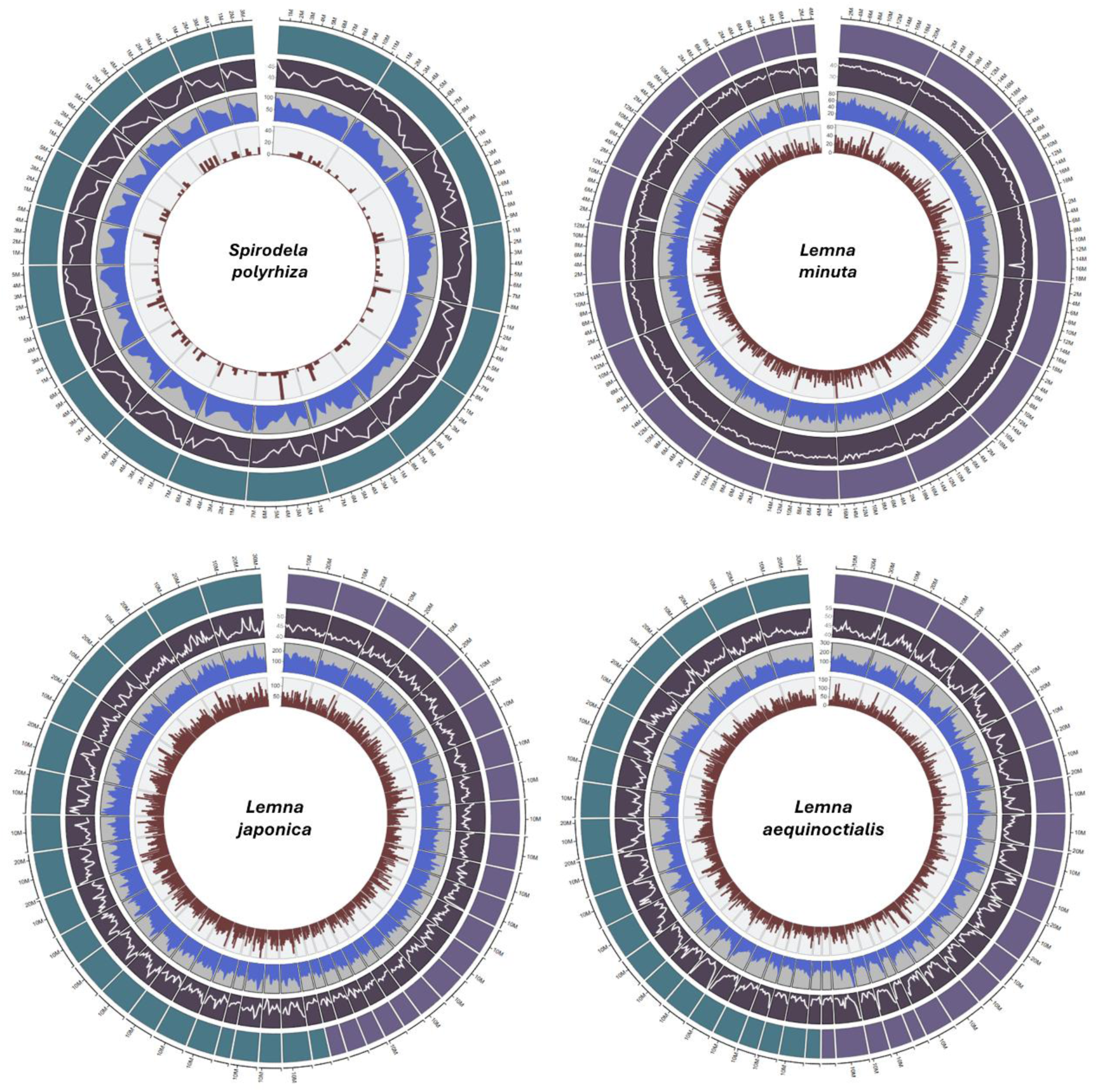
Genome composition and annotation across four duckweed species. Circos plots illustrating key genomic features along the chromosomes of *Spirodela polyrhiza*, *Lemna minuta*, *L. japonica*, and *L. aequinoctialis* ×. Tracks are shown from outermost to innermost as follows: (i) chromosome length; (ii) chromosome ideograms numbered sequentially; (iii) GC content; (iv) gene density; and (v) transposable element (TE) density. In *L. japonica*, the two subgenomes are distinguished by colour, with subgenome M (*L. minor*-derived) shown on the right and subgenome T (*L. turionifera*-derived) on the left. In *L. aequinoctialis* ×, subgenome A (*L. aequinoctialis*-derived) is shown on the left and subgenome X (unknown parental origin) on the right, with distinct colours reflecting their different ancestries. Genomic features were calculated using sliding windows of 500 kb for *L. minuta* and *S. polyrhiza*, and 1 Mb for *L. japonica* and *L. aequinoctialis* ×.

We next compared the abundance of annotated InterProScan terms across species. Pentatricopeptide repeats (PPR) (IPR002885) were the most abundant across all species, with counts ranging from 2,942 in *L. minuta* to 5,987 in *L. japonica*, and 3,399 and 5,565 in *S. polyrhiza* and *L. aequinoctialis* ×, respectively. Protein kinase domains (IPR000719) were also highly represented, with 531 in *S. polyrhiza* and 439 in *L. minuta*, increasing to 799 and 794 in *L. japonica* and *L. aequinoctialis* ×.

Despite its smaller genome size, *S. polyrhiza* showed higher representation of several metabolism-associated domains compared with *L. minuta*, including Cytochrome P450 (CYP450; 198 vs 155), UDP-glucuronosyl/glucosyltransferase (UGT; 68 vs 44), and haem peroxidase (70 vs 57). In contrast, defence-related domains such as NB-ARC (IPR002182) were more abundant in *L. minuta* (149) than in *S. polyrhiza* (73), while antimicrobial protein (MiAMP) families were also enriched in *L. minuta* (91) relative to *S. polyrhiza* (66) and *L. aequinoctialis* × (99).

Transcription factor-associated domains further highlighted these differences: WRKY (64 vs 59), SANT/Myb (201 vs 158), and bHLH (77 vs 49) domains were consistently more abundant in *S. polyrhiza* than in *L. minuta*. In the larger and hybrid species *L. aequinoctialis* × (99, 336, and 105) and *L. japonica* (102, 330, and 111), domain counts are higher overall and show only limited variation between the two species, likely reflecting their shared genome expansion associated with hybridisation. In contrast, despite its smaller genome, *S. polyrhiza* consistently shows higher domain counts than *L. minuta*, which may indicate a relative expansion of transcription factor-associated domains (Figure 4c).

**Figure 4.**
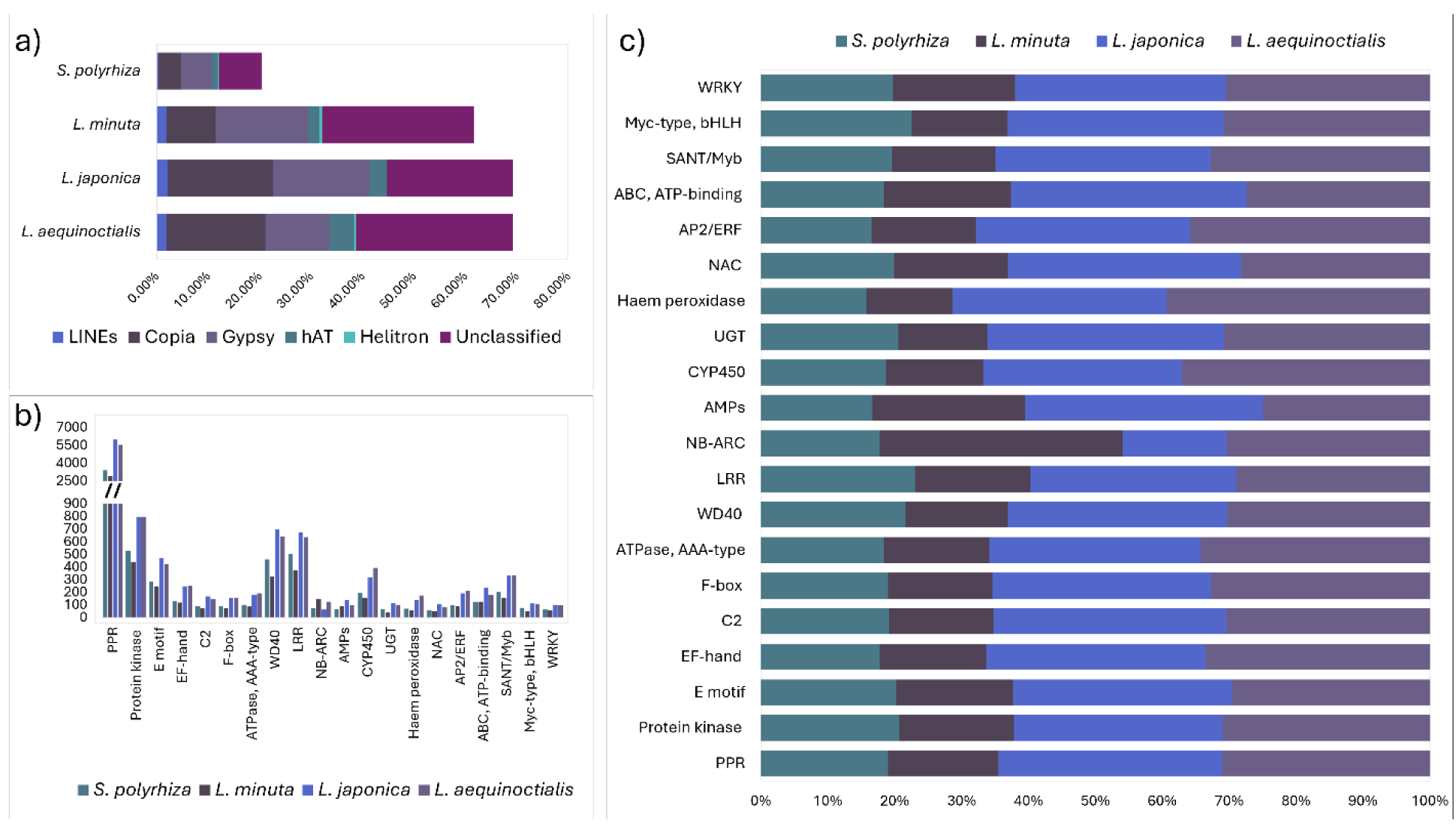
Transposable element composition and functional annotation across four duckweed species. a) Stacked bar chart showing the distribution of transposable element and repeat classes identified using RepeatModeler across the four species. b) Clustered column chart displaying the most abundant InterProScan annotation terms, including domains, repeats, and protein families, in each species. c) Percentage stacked bar chart comparing the relative abundance of InterProScan terms across *S. polyrhiza*, *L. minuta*, *L. japonica*, and *L. aequinoctialis* ×. Raw data are provided in Files S2 and S3.

### Gene Family Expansion and Reduction

Gene family evolution across duckweeds revealed distinct lineage-specific patterns (Figure 5a). The estimated global birth–death parameter (λ = 1.79) approached the upper bound of the model, indicating a high rate of gene family turnover across the phylogeny. At the Lemnaceae node, gene family contractions (−1,889) clearly exceeded expansions (+316), a trend that was also evident in the aquatic lineage including duckweeds and *Z. marina* (+8 / −621). Within *Spirodela*, contractions dominated in both *S. polyrhiza* (+718 / −1,127) and *S. intermedia* (+375 / −484), as well as at the *Spirodela* node (+260 / −1,031). In contrast, the *Lemna* lineage showed an overall excess of gene family expansions at the genus node (+2,591 / −794). This expansion pattern was particularly pronounced in hybrid *L. japonica* (+5,754 / −1,646) and *L. aequinoctialis* × (+5,488 / −834). Expansion-dominated patterns were also observed in *L. turionifera* (+1,790 / −766), whereas *L. minor* displayed a more balanced pattern with slightly higher contractions (+1,133 / −1,372). In contrast, *L. minuta* exhibited a strong excess of contractions (+372 / −4,124) (File S10).

**Figure 5.**
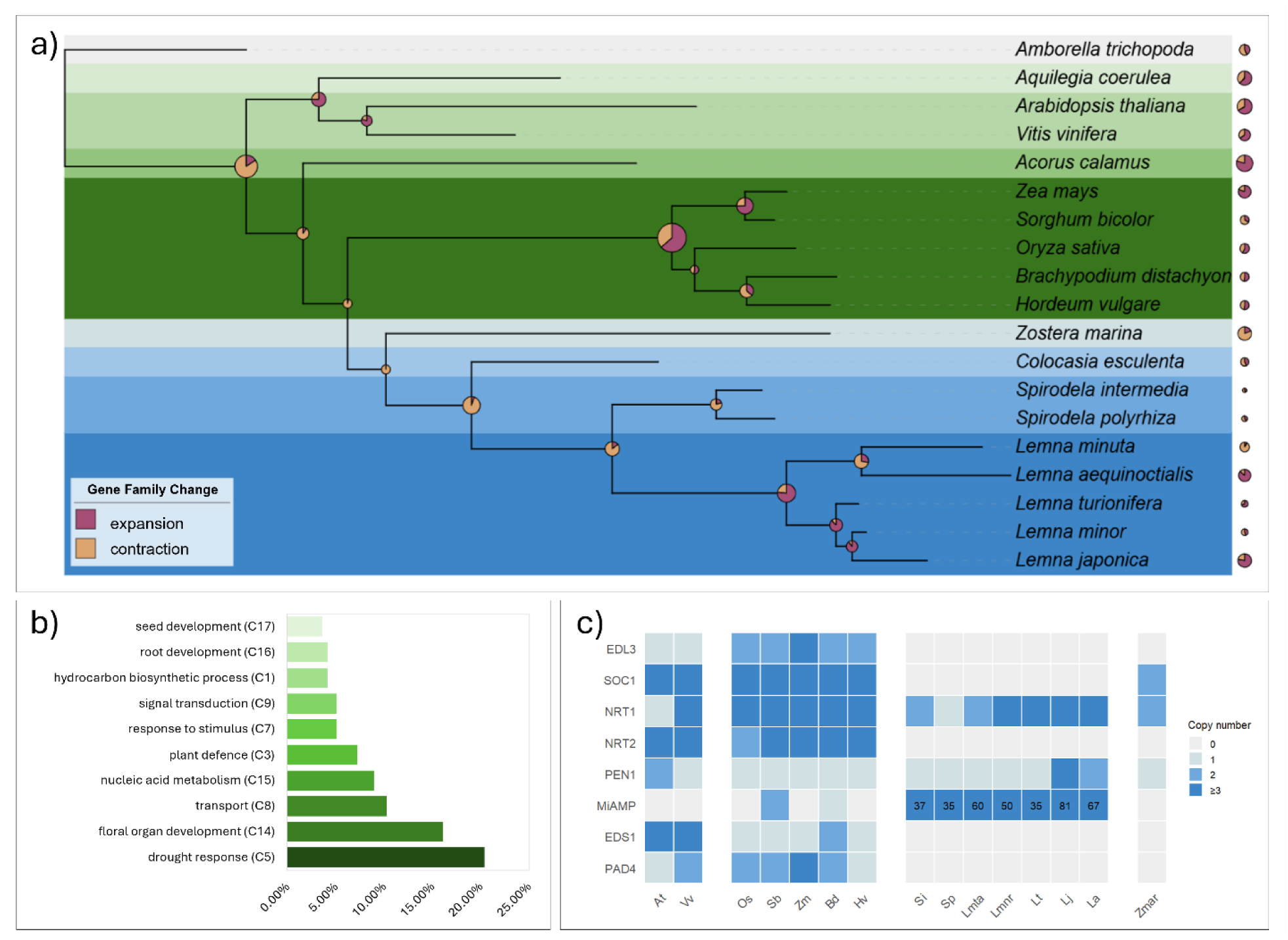
Gene family evolution and functional clustering of missing orthogroups in duckweeds. a) Phylogenetic tree inferred using OrthoFinder, illustrating gene family expansions and contractions across selected terrestrial and aquatic plant species, including duckweeds. Gene family dynamics were determined using CAFE5. Coloured circles at each node indicate gene family changes, with purple representing expansion and orange representing contraction; circle size is proportional to the number of gene families affected. The complete phylogenetic tree with node labels and corresponding expansion and contraction values is provided in Figure S7 and File S10. b) Bar chart showing the number of missing orthogroups per functional cluster, based on GO enrichment analysis performed using DAVID. Missing orthogroups were defined as those present in *A. thaliana* and *O. sativa* but absent across all duckweed species examined. Complete annotation for all clusters is provided in File S5. c) Tile plot displaying the presence and absence of selected orthogroups of biological interest across species, including proteins associated with plant defence: PENETRATION (PEN), ENHANCED DISEASE SUSCEPTIBILITY 1 (EDS1), and PHYTOALEXIN DEFICIENT 4 (PAD4), and antimicrobial proteins (MiAMP); nitrate transport: nitrate transporters NRT1 and NRT2; and ABA signalling and flowering: EID1-like 3 (EDL3) and SUPPRESSOR OF OVEREXPRESSION OF CONSTANS 1 (SOC1).

Absent orthogroups (present in *Arabidopsis thaliana* and *Oryza sativa* but absent from all duckweed species) were grouped into 15 enriched Gene Ontology clusters (File S5). The largest cluster (Cluster 5) was dominated by drought response and ABA-related terms, including response to water deprivation (n = 21), response to water (n = 21), cellular response to oxygen-containing compounds (n = 25), and signal transduction (n = 26). All duckweed species lacked EID1-like 3 (EDL3; OG0014185), associated with ABA signalling (Figure 5c) but *Colocasia esculenta* contains 2 EDL3 in same orthogroup (File S9).

The second largest cluster (Cluster 14) comprised developmental processes, including system development (n = 34), post-embryonic development (n = 26), reproductive system development (n = 21), and shoot system development (n = 14). SUPPRESSOR OF OVEREXPRESSION OF CONSTANS 1 (SOC1) (OG0001658) was absent in all duckweed species but present in *Zostera marina* (Figure 5c). Root- and seed-related processes were also represented in Cluster 16 (root development, n = 9) and Cluster 17 (seed development, n = 8).

Transport-related functions were grouped in Cluster 8, including nitrogen compound transport (n = 24) and protein transport (n = 15). The high-affinity nitrate transporter NRT2 (OG0008317) was absent in all duckweeds and *Z. marina*, whereas NRT1 (OG0003002) was retained across all species (Figure 5c). Cluster 15 included nucleic acid metabolic processes (n = 22–25 terms).

Additional clusters were associated with responses to stimuli, defence, and signalling (Clusters 3, 7, and 9). Cluster 3 included defence-related processes such as defence response to other organism (n = 16), response to salicylic acid (n = 8), and hypersensitive response-related terms (n = 6). Key immunity-related genes, including PAD4 (OG0012145), EDS1 (OG0012594), and CC-NBS-LRR (OG0006349), were absent across all duckweed species and *Z. marina*.

In contrast, antimicrobial peptide genes (MiAMP; OG0000103) were expanded in duckweeds, with copy numbers of 35 in *S. polyrhiza*, 37 in *S. intermedia*, 60 in *Lemna minuta*, 50 in *L. minor*, 35 in *L. turionifera*, 81 in *L. japonica*, and 67 in *L. aequinoctialis* × (Figure 5c). PEN proteins (OG0012030) were retained across duckweeds and *Z. marina*, indicating partial conservation of defence-related components. Moreover, selected orthogroups of biological interest were confirmed through cross-referencing OrthoFinder identifiers with EggNOG and InterProScan annotations and validated by querying *A. thaliana* gene identifiers against the filtered orthogroup output (File S11).

### Pathway Mapping for Secondary Metabolite Biosynthesis

Annotation of secondary-metabolite biosynthesis pathways revealed no contraction comparable to that observed for organ-development and stress-response gene families. Instead, flavonoid-related pathways (flavonoid, anthocyanin, flavone, and flavonol biosynthesis) showed high gene counts across all duckweed genomes relative to land-plant references (Table 2). Flavonoid biosynthesis genes ranged from 42 to 89 across duckweeds, compared with 31 in *A. thaliana* and 40 in *O. sativa*. Carotenoid biosynthesis genes were also abundant, with 55 copies in *S. polyrhiza*, 43 in *L. minuta*, 86 in *L. japonica*, and 114 in *L. aequinoctialis ×*, exceeding *A. thaliana* (32) and comparable to or higher than *O. sativa* (40).

**Table 2.**
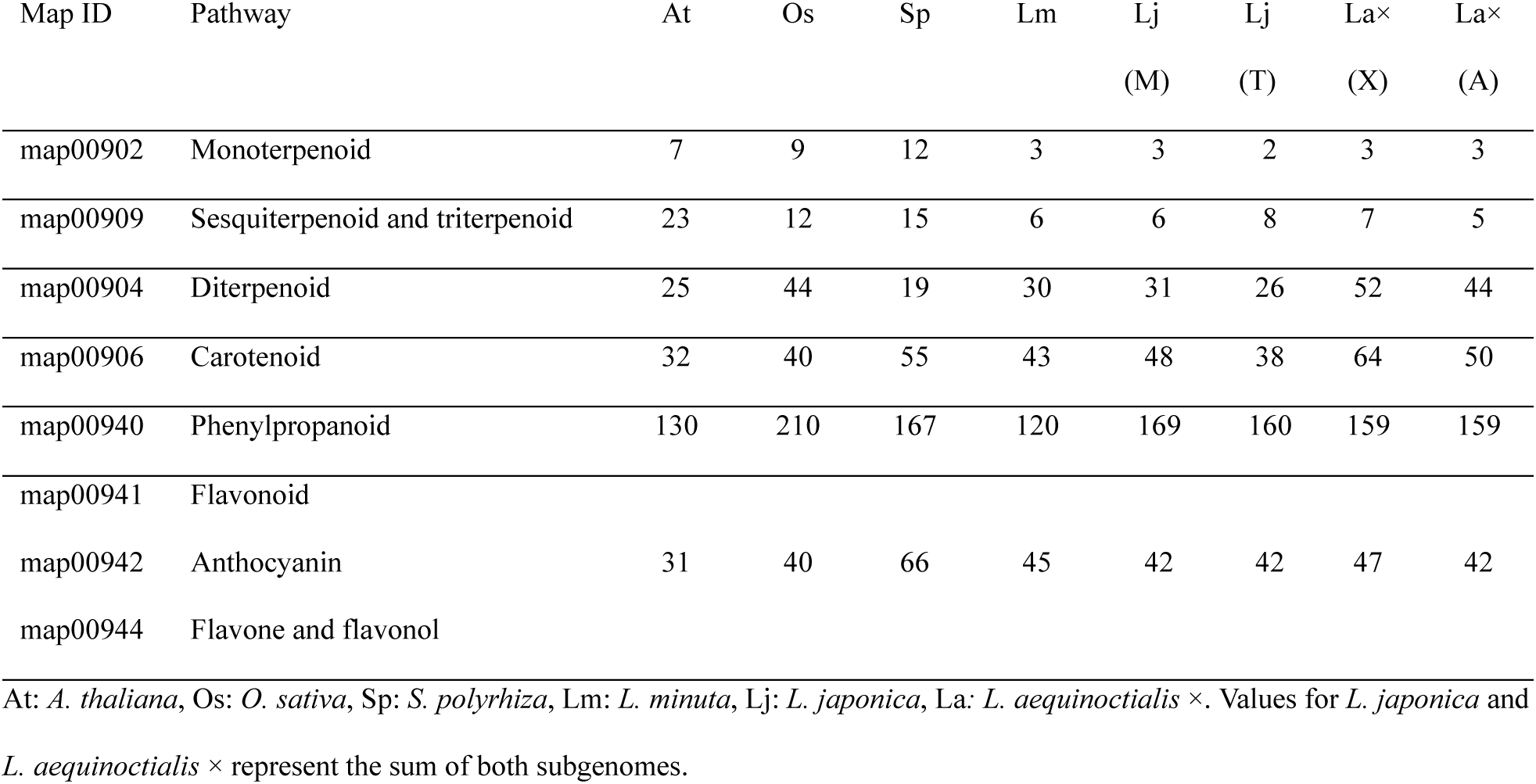
Gene counts associated with secondary metabolite biosynthesis pathways in duckweeds compared with *A. thaliana* and *O. sativa*, based on KEGG and EggNOG annotations.

Across multiple pathways, *L. minuta* consistently showed lower gene counts relative to other duckweeds, including carotenoid (43 genes) and phenylpropanoid biosynthesis (120 genes), compared with 167 in *S. polyrhiza*. Diterpenoid biosynthesis genes ranged from 30 to 96 across duckweeds, compared with 25 in *A. thaliana* and 44 in *O. sativa*. Sesquiterpenoid and triterpenoid biosynthesis genes ranged from 5 to 14 copies across duckweeds, compared with 23 in *A. thaliana* and 12 in *O. sativa*, whilst monoterpenoid biosynthesis genes remained lower across all duckweed species (3-6 copies). In the phenylpropanoid biosynthesis pathway, *L. minuta* (120) contained slightly fewer genes than the other species examined. Although *S. polyrhiza* (167) contained more mapped genes than *L. minuta*, the total remained lower than in *O. sativa* (210). Hybridisation appears to have contributed to pathway expansion in *L. japonica* (M: 169, T:160) and *L. aequinoctialis* × (A:159, X:159); however, when individual subgenomes were considered separately, gene counts in the phenylpropanoid pathway remained lower than in *O. sativa*.

## Discussion

Genome evolution in duckweeds following the return to water is not passive erosion but an environmentally coherent programme: costly immunity components that impose little benefit under low pathogen pressure are selectively lost, high-affinity nutrient transporters become dispensable in nitrogen-rich water, organ-development regulators are redundant without roots and flowers [3], yet secondary-metabolite pathways [4] or antimicrobial peptides that provide chemical defence and abiotic buffering are preserved and, in some lineages, expanded [25]. Our chromosome-scale assemblies illuminate this programme through three key discoveries: an unresolved hybrid lineage with a post-hybridisation translocation, convergent losses shared with seagrasses (*Z. marina*), and differential retention of biosynthetic capacity.

Interspecific hybridisation is a major driver of genetic variation, but most interspecific hybrids face rapid extinction [26]. Nevertheless, hybridisation appears relatively common within duckweeds; *L. japonica*, whose origin from *L. minor* and *L. turionifera* was confirmed here through Omni-C contact maps and genome-wide synteny analysis (Figures 1c and 2a), corroborating previous Hi-C-based evidence [3]. Other documented hybrids include *L.* × *mediterranea*, derived from *L. gibba* and *L. minor*, and *L.* × *aoukikusa*, derived from *L. aequinoctialis* and *L. perpusilla* [10,13]. The strain identified as *L. aequinoctialis* D3 displayed hybrid genome signals in Omni-C contact maps, with BUSCO results dominated by duplicated hits, supporting a hybrid origin (Figure 1d, Table 1). Attempts to resolve parental identity using rDNA sequences were unsuccessful owing to identical sequences across known hybrid lineages (File S8), and whole-genome synteny analysis was not possible given the absence of chromosome-scale reference genomes for candidate parental species, including *L.* × *aoukikusa* and *L. perpusilla* [10]. This assembly is therefore interpreted as comprising an *L. aequinoctialis*-derived subgenome (A) and an unresolved parental subgenome (X). SOC1, a central regulator of the vegetative-to-reproductive transition [27], is absent across most duckweed species [3,28], although fertile *L. aequinoctialis* strains represent a notable exception, retaining SOC1 expression alongside other MADS-box transcription factors [29]. Its absence in *L. aequinoctialis* × may reflect loss during or following hybridisation, potentially contributing to the reproductive sterility observed in related backcross hybrid lineages [10].

Beyond genome size and hybrid origin, duckweed species also differ markedly in genome composition [7]. TE content was lowest in *S. polyrhiza* (20.91%), consistent with its highly compact genome and previously reported high gene density [3,4]. In contrast, the *Lemna* species exhibited substantially higher repeat fractions, with *L. aequinoctialis* (73.20%) and *L. japonica* (72.50%) showing the most repeat-rich genomes among those analysed here. *L. minuta* displayed a slightly lower but still substantial TE content (65.47%), in close agreement with the ∼58.2% previously predicted for this species [23], and notable given that it harbours the smallest genome (∼308 Mb) among the *Lemna* species examined. *S. polyrhiza* also showed a distinct TE composition, with LTR/Gypsy elements (6.02%) more abundant than LTR/Copia elements (4.39%) (Figure 3, File S2), consistent with previous genome annotations [3,4]. A Gypsy/Copia ratio of ∼3.5 has previously been reported for *S. polyrhiza* [28], whereas we observed a ratio of ∼1.4 — a difference that likely reflects variation in repeat annotation methodology or accession-specific differences in Copia element content rather than a genuine biological shift.

Chromosomal rearrangements represent another major force in genome evolution, generating structural diversity through micro- and macro-scale genome shuffling [30]. Although their direct fitness consequences remain unpredictable, chromosomal rearrangements represent a significant source of structural variation that can shape genome evolution and contribute to lineage divergence [31,32]. Such rearrangements have been previously reported in duckweeds, including translocations, inversions, and chromosomal fissions [3,8,33]. Here, we identified a ∼3.5 Mb reciprocal translocation between chromosome 12 of subgenome X and chromosome 17 of subgenome A in *L. aequinoctialis* × (Figure 2, Figure S2), localised at the telomeric regions of both chromosomes, consistent with the tendency of macro-rearrangements to cluster near telomeres and centromeres [34]. Although such structural changes are frequently associated with polyploidy or interspecific hybridisation[35], whether this translocation contributes to genome divergence or fitness differences in *L. aequinoctialis* × remains to be determined.

PPR domains (IPR002885) were the most abundant InterProScan annotation terms across all four species (Figure 4), with EggNOG identifying 405, 402, 800, and 1,083 PPR-containing proteins in *S. polyrhiza*, *L. minuta*, *L. japonica*, and *L. aequinoctialis* ×, respectively (File S3), the higher counts in hybrid species likely reflecting subgenome dosage. PPR proteins constitute one of the largest protein families in land plants (441 and 655 members in *A. thaliana* and *O. sativa*, respectively), where they function in organellar RNA metabolism, including processing, modification, and translation [36,37]. Their expansion is considered a hallmark of terrestrialisation, as green algae harbour far fewer members (*Chlamydomonas reinhardtii*, 14; *Volvox carteri*, 10) [38,39]. Despite pervasive gene family loss in duckweeds, PPR proteins remain well represented at land plant levels, with comparable numbers (579) reported in *L. punctata* [4]. This trajectory, from minimal algal repertoires through expansion during terrestrialisation to retention in secondarily aquatic duckweeds, suggests that the organellar RNA editing machinery, once established, persists independently of habitat and may have contributed to adaptation across multiple evolutionary transitions.

Duckweeds are well known for their extreme morphological reduction, reflected at the genomic level by widespread gene family contraction and loss [2]. Losses in gene families associated with DNA/RNA methylation, signal transduction, and organ development have been reported previously [3,4], and our orthogroup-based analysis broadly corroborates these findings (Figure 5b). The largest contraction cluster was enriched for drought stress response terms (File S5 and S9), consistent with an aquatic lifestyle in which drought stress is absent as a selective pressure [4,40]. Among completely lost orthogroups, EDL3 (OG0014185), an F-box protein mediating ABA-dependent drought responses [41], was absent across all assembled duckweed genomes and in *S. intermedia*, *L. minor*, and *L. turionifera* (File S9).

Beyond previously characterised losses, we identified missing orthogroups with functional implications less explored in duckweed genomics (Figure 5c). The high-affinity nitrate transporter NRT2 (OG0008317) was absent across all duckweed genomes and in *Z. marina*, but retained in *C. esculenta* (File S9), suggesting convergent loss of high-affinity nitrate uptake in fully aquatic but not semi-aquatic lineages. NRT2 and NRT1 mediate high-affinity and low-affinity NO₃⁻ uptake, respectively, and constitute the two major nitrate transporter families in plants [42]. NRT1 (OG0003002) was retained across all species examined, indicating preserved basal nitrate transport capacity and pointing to a functional shift toward low-affinity uptake consistent with nitrogen-rich aquatic environments [43]. More broadly, reduction of transport-associated gene families likely reflects the simplified duckweed body plan, which lacks differentiated organ systems and the associated requirements for long-distance solute transport.

Alongside the reduction of abiotic stress and transport-related gene families, duckweed genomes exhibit a fundamental remodelling of immunity reflecting the distinct biotic pressures of aquatic environments Plant-pathogen interactions have driven plant evolution and contributed to land colonisation [44], with ETI evolving from the ancestral PTI system through R gene diversification and EDS1/PAD4 acting as core regulators linking ETI to SAR signal transduction [45,46]. Both EDS1 and PAD4, alongside CC-NBS-LRR proteins, were absent across all duckweed species and in *Z. marina* (Figure 5c, File S9), corroborating and extending previous reports in *L. punctata*, *S. polyrhiza*, and *W. australiana* [25].

Despite this remodelling, duckweeds are not defenceless. Within the PTI layer, PEN1, a syntaxin (SYP121) central to preinvasive immunity [47], was retained across duckweeds and *Z. marina*, and MiAMPs were substantially expanded in duckweeds but absent in *Z. marina* and most of the land plants including *A. thaliana* and *O. sativa* (Figure 5c, File S9), reflecting lineage-specific elaboration of broad-spectrum defence. Together with expanded secondary metabolite biosynthesis (Table 2), these PTI components represent a preformed chemical barrier against pathogen ingress [48], reflecting a shift from costly ETI-based immunity toward ancestral broad-spectrum resistance sufficient for their ecological context.

This pattern is consistent with relaxed Red Queen coevolutionary dynamics [49,50]:Terrestrial environments generally harbour a higher microbial load than aquatic surface habitats [51], likely imposing less selective pressure to maintain costly ETI-based immunity following the return to water [52]. Interestingly, this model does not fully apply to *L. minuta*, one of the most invasive duckweed species [6]. Despite its relatively small genome, *L. minuta* showed greater NB-ARC domain (149) abundance than other *Lemna* species examined (Figure 4c), representing an unexpected deviation from the general pattern of immune reduction. This expansion may reflect selection for broader pathogen recognition capacity during range expansion into novel environments harbouring unfamiliar microbial communities, consistent with the ’enemy release followed by novel enemy encounter’ model of invasive plant success [53]

Notably, despite its compact genome, *S. polyrhiza* was richer than *L. minuta* in both CYP450 (IPR001128) and UGT (IPR002213) families (Figure 4b). CYP450s participate in secondary metabolite biosynthesis and metabolic diversification [54,55], whilst UGTs catalyse glycosylation of diverse substrates, including flavonoids [56]. This enrichment is particularly striking because domain family size generally scales with genome size across the species examined here (Figure 4c), making the pattern in *S. polyrhiza* an exception worth highlighting. The abundance of CYP450s and UGTs also correlated with the number of genes assigned to secondary metabolic pathways relative to *O. sativa* and *A. thaliana* (Table 2). Despite the broad gene family loss characteristic of duckweeds, genes associated with flavonoid, anthocyanin, flavone, and flavonol biosynthesis pathways appear largely retained (Table 2). *S. polyrhiza*, *L. japonica*, and *L. aequinoctialis* × showed the greatest expansion of genes in these pathways among the species examined, consistent with the reported enrichment of flavonoid biosynthesis genes in *L. punctata* relative to *A. thaliana* and *O. sativa* [4]. *L. minuta*, by contrast, showed comparatively limited expansion of secondary metabolite biosynthesis genes (Table 2). Taken together, these findings suggest that the capacity for secondary metabolite biosynthesis, particularly flavonoid production, has been preferentially retained in duckweeds despite overall genome compaction.

## Conclusion

Overall, our work provides chromosome-scale genomic resources for four duckweed species, reveals the unresolved hybrid nature of *L. aequinoctialis* ×, and demonstrates that the return to an aquatic lifestyle has driven a coherent, convergent programme of genome remodelling shared across independent aquatic lineages. These findings advance our understanding of how habitat transitions shape genome architecture and remodel key functional pathways and offer a high-quality genomic foundation for future research in plant evolutionary biology and biotechnology.

## Methods

### Plant material

Four duckweed accessions (D1, D2, D3, and D4) were obtained from Aplantex Inc., Québec (QC), Canada, and taxonomically identified as *L. minuta* (D1), *L. japonica* (D2), *L. aequinoctialis* (D3), and *S. polyrhiza* (D4). Accessions were cultivated in half-strength Hoagland medium (0.65 g/L) supplemented with 0.5% MES, following surface sterilisation with 70% ethanol and 5% bleach, and rinsed three times with sterile water.

### High Molecular Weight (HMW) DNA extraction and sequencing

Genomic DNA was extracted from *S. polyrhiza*, *L. minuta*, *L. japonica*, and *L. aequinoctialis* × using a CTAB-based method [57,58] following grinding of 500-1,000 mg of frond tissue in liquid nitrogen. Following quantification, high molecular weight DNA (>2 µg) was used for library preparation according to the manufacturer’s protocol and sequenced on the PacBio HiFi platform. Extracted DNA (∼300 ng) was additionally used for short-read sequencing on the Element AVITI™ platform. For chromatin conformation capture, 300 mg of young fronds were ground in liquid nitrogen and processed using the Dovetail Omni-C Kit (Dovetail Genomics, Scotts Valley, CA, USA) according to the manufacturer’s instructions.

### Genome assembly

The genome assembly procedure was tailored to the available dataset for each of the four species. PacBio HiFi long reads were used to construct draft genome assemblies using Flye v2.9.5 [59] or hifiasm 0.25.0-r726 [60] with various parameters sets. Based on QUAST curves (version 4.6.1) [61], the draft assembly that had the most assembled nucleotides within the fewest contigs was selected for downstream assembly steps. Duplicate contigs were subsequently removed using the purge_dups (version 1.2.6) [62] pipeline. To remove potential contaminant sequences, assembled contigs were taxonomically classified using Kraken 2.1.3 [63,64], FCS-GX 0.5.0 (FCS-GX database version 1; build date: 2023-01-24) [65] and/or the BlobToolKit procedure (version 4.4.4) [66]. Suspected contaminant contigs hereby identified were filtered out using *seqkit* grep.

Omni-C data was then used for scaffolding using YaHS 1.1, after processing the data using the pairtools (version 1.1.2) workflow f [67,68]. Omni-C data was then used to generate chromatin contact matrices, which were manually curated using PretextView [69]. Contigs that could not be incorporated into the main scaffolds based on the expected chromosome number, or could not be confidently positioned, were designated as unplaced contigs.

Following generation of the final assembly in FASTA format using the sanger-tol workflow [67], unplaced contigs were excluded from downstream analyses including gene prediction and genome annotation. For *Spirodela polyrhiza*, the available AVITI short reads were used to polish the draft assembly by first mapping them to the main scaffolds using minimap2 [70], then running Hapo-G 1.3.8 [71]. Subgenome phasing of *L. japonica* and *L. aequinoctialis* × was performed using Subphaser 1.2.7 and visualised using D-GENIES [72,73]. For *L. minuta*, *L. aequinoctialis* and *L. japonica*, organelles were assembled from reads using oatk 1.0 [74] with the embryophyta lineage dataset. Assembly quality was evaluated using merqury 1.3 and compleasm 0.2.7 for assembly statistics and genome completeness, respectively [61,75]. A schematic overview of the assembly workflow for each species is provided in Figures S3–S6.

### Synteny analysis

In addition to the genomes assembled in this study, previously published genome assemblies were included in synteny and downstream analyses: *Spirodela intermedia* 8410 obtained from NCBI (GCA_902729315.2)[8], and *Lemna minor* 7210 and *Lemna turionifera* 9434 obtained from lemna.org[3]. Synteny analysis was performed using DEEPSPACE to visualise collinearity among duckweed species, using chromosome-scale genome assemblies in FASTA format as input[76]. Default parameters were used for parsing and scanning with minimap2 and MCScanX, except for the parameter *minChrLen*, which was set to 1,000,000 bp to account for the relatively small genome sizes and shorter chromosome lengths observed in *S. polyrhiza* [70,77]. Raw riparian plots are provided in File S6 for both visualisation options: (i) chromosomes scaled to the same x-range across genomes, and (ii) chromosomes scaled according to their physical sizes. Chromosomal labels were removed from the figures to improve visual clarity. As a hybridisation signal was detected during the manual curation step for *L. aequinoctialis* ×, an additional intra-genomic synteny analysis was performed between the two subgenomes. The original output of this analysis is provided in File S7.

### Genome annotation

De novo transposable element (TE) annotation was performed using RepeatModeler 2.0.1 [78] followed by RepeatMasker v4.1.1 [79] on the assembled genomes. Gene prediction was performed using AUGUSTUS for *ab initio* gene prediction [80], with the parameters *--strand* both, *--genemodel* complete, *--gff3*, *--protein*, *--codingseq*, *--cds*, and *--softmasking*.

Predicted protein sequences were functionally annotated using InterProScan v5 (database release 5.59-91.0), including Pfam domain annotations [81,82]. InterProScan and Pfam identifiers were extracted and processed using pandas for clustering and counting of protein domains, families, and repeat-associated motifs. Predicted protein sequences were additionally annotated using EggNOG-mapper with the Viridiplantae database and the DIAMOND search mode (*-m* diamond) [83,84].

### Ortholog identification and detection of missing genes

To identify orthologous gene groups across aquatic and terrestrial plant lineages, proteomes from several species were included in the analysis These comprised the ancestral monocot *Acorus calamus* (Eurasian sweet-flag) from NCBI (GCA_030737845.1) [85]; the basal eudicots *Aquilegia coerulea* (GCA_002738505.1) [86]; the semi-aquatic Araceae species *C. esculenta* (GCA_053894955.1), which belongs to the same family as duckweeds; and the aquatic angiosperm *Zostera marina* (GCA_001185155.1). Representative monocot crop species were also included: *Brachypodium distachyon* (GCF_000005505.3), *Hordeum vulgare* (GCF_904849725.1), *Oryza sativa* (GCF_034140825.1), *Sorghum bicolor* (GCF_000003195.3), and *Zea mays* (GCF_902167145.1). Two eudicot species, *A. thaliana* (GCF_000001735.4) and *Vitis vinifera* (GCF_030704535.1), were additionally included. These datasets were analysed together with the duckweed genomes assembled in this study, as well as previously published duckweed genomes used in the synteny analyses [3,8].

Protein sequences from all species were analysed using OrthoFinder to identify orthologous gene groups [87], with the parameters *-S* diamond, *-T* iqtree, and *-A* mafft. Orthogroup counts and duplication events were further processed using NumPy, pandas, and Excel for filtering, comparative analyses, and extraction of gene identifiers. Gene family expansion and contraction across species and phylogenetic nodes were estimated using CAFE5 [88].

To detect potentially missing genes in duckweed genomes, hierarchical orthogroups showing zero counts across all duckweed species but present in both *A. thaliana* and *O. sativa* were extracted. These orthogroups were subsequently annotated using the DAVID functional annotation tool for Gene Ontology (GO) enrichment, combining gene identifiers from *A. thaliana* and *O. sativa* [89]. The phylogenetic tree was visualised using iTOL [90] with the SpeciesTree output generated by OrthoFinder, and further annotated with pie charts representing gene family expansion and contraction patterns derived from the CAFE5 analysis.

Selected orthogroups of biological interest were identified through a combination of approaches. First, orthogroups flagged within functionally enriched clusters following DAVID GO annotation were manually inspected, and genes of known relevance to plant adaptation, development, and immunity were prioritised for further examination. Second, *A. thaliana* or *O. sativa* gene identifiers associated with biological processes of interest were used to query the filtered orthogroup output directly, allowing targeted retrieval of specific orthogroups absent across all duckweed species. Third, orthogroups showing notable expansion or contraction in CAFE5 results were cross-referenced with EggNOG and InterProScan annotations to assign putative functions. Gene identities were confirmed by cross-referencing orthogroup annotations from both EggNOG-mapper and InterProScan, and selected orthogroups are reported with their corresponding OrthoFinder identifiers (File S9).

### Pathway annotation and comparison

Secondary metabolite biosynthesis pathways were identified using KEGG pathway map IDs obtained from EggNOG annotations and verified using KEGG Mapper [91]. The following pathways were analysed: monoterpenoid biosynthesis (*map00902*), sesquiterpenoid and triterpenoid biosynthesis (*map00909*), diterpenoid biosynthesis (*map00904*), carotenoid biosynthesis (*map00906*), phenylpropanoid biosynthesis (*map00940*), flavonoid biosynthesis (*map00941*), anthocyanin biosynthesis (*map00942*), and flavone and flavonol biosynthesis (*map00944*).

Genes associated with these pathways were extracted from EggNOG annotation results for *L. minuta*, *L. aequinoctialis* ×, *L. japonica*, and *S. polyrhiza* following filtering by map ID. Gene identifiers were subsequently counted for each pathway. Given the substantial overlap among genes associated with flavonoid-related pathways (flavonoid, anthocyanin, and flavone/flavonol biosynthesis), genes from these pathways were combined and reported as a single group. For hybrid species (*L. aequinoctialis* × and *L. japonica*), genes from both subgenomes were considered to better represent total gene content and avoid underestimation due to subgenome partitioning. For comparison, gene counts for *A. thaliana* and *O. sativa*, representing well-annotated reference species for eudicots and monocots in the KEGG database, were obtained using the same pathway IDs and compared with duckweed species.

## Supplementary information

**Figure S1.** GenomeScope profiles generated from PacBio HiFi reads for each species.

**Figure S2.** Omni-C contact map illustrating the reciprocal translocation between chromosomes 12X and 17A in *L. aequinoctialis ×*.

**Figure S3.** Genome assembly pipeline for *S. polyrhiza*.

**Figure S4.** Genome assembly pipeline for *L. minuta*.

**Figure S5.** Genome assembly pipeline for *L. aequinoctialis ×*.

**Figure S6.** Genome assembly pipeline for *L. japonica*.

**Figure S7.** Rooted phylogenetic tree generated by OrthoFinder with node and species identifiers, provided for interpretation of CAFE5 results.

**File S1.** Assembly statistics for each genome, including contiguity and completeness metrics from QUAST and BUSCO.

**File S2.** Transposable element annotation generated with RepeatModeler, including element counts and classifications for each species.

**File S3.** InterProScan annotation results including Pfam and InterPro domain identifiers and descriptions for each species.

**File S4**. EggNOG functional annotation including KEGG pathway assignments for each species.

**File S5.** GO term annotation of contracted gene clusters with gene counts, generated using DAVID.

**File S6.** Riparian plots for genome-wide synteny analysis generated with DEEPSPACE among duckweed species, including scaled and unscaled versions with chromosome identifiers.

**File S7.** Riparian plots for synteny analysis between *L. aequinoctialis* × subgenomes generated with DEEPSPACE, including scaled and unscaled versions with chromosome identifiers.

**File S8.** Genetic similarity matrix of rDNA sequences across duckweed species, used to investigate the parental origin of L. aequinoctialis ×.

**File S9.** OrthoFinder output tables including orthologous gene identifiers and counts across species.

**File S10.** CAFE5 results detailing contracted and expanded gene family numbers at internal nodes and terminal species in the phylogenetic tree.

**File S11.** NCBI gene and protein identifiers of selected orthogroups across available species, cross-referenced with InterProScan and EggNOG annotations in duckweeds.

## Data Availability

Genome assemblies and annotation files are available upon request at Zenodo (ID: 19823072) https://doi.org/10.5281/zenodo.19823072

## Author contributions

MIT: methodology, data curation, analysis, annotation, writing draft, reviewing and editing manuscript; BB: sequence data generation; EN: genome assembly; LL: genome assembly; MS: funding acquisition; DT: planning and designing, project administration, funding acquisition, reviewing and editing manuscript

## Acknowledgments

This work was supported by [ALLRP 585444 - 23] from Natural Sciences and Engineering Research Council of Canada (NSERC) Alliance Grant and Consortium de recherche et innovations en bioprocédés industriels au Québec (CRIBIQ) to DT. The authors thank IBIS’s Genomic Platform for PacBio HiFi and AVITI sequencing, and Aplantex Inc. for providing duckweed accessions.

## Ethics approval and consent to participate

Ethics approval is not applicable.

