## Supplementary figures and images for "Chromosome-scale genome assemblies of duckweeds provide insights into genomic plasticity, aquatic adaptation and morphological reduction"

### Supplementary Figure 1

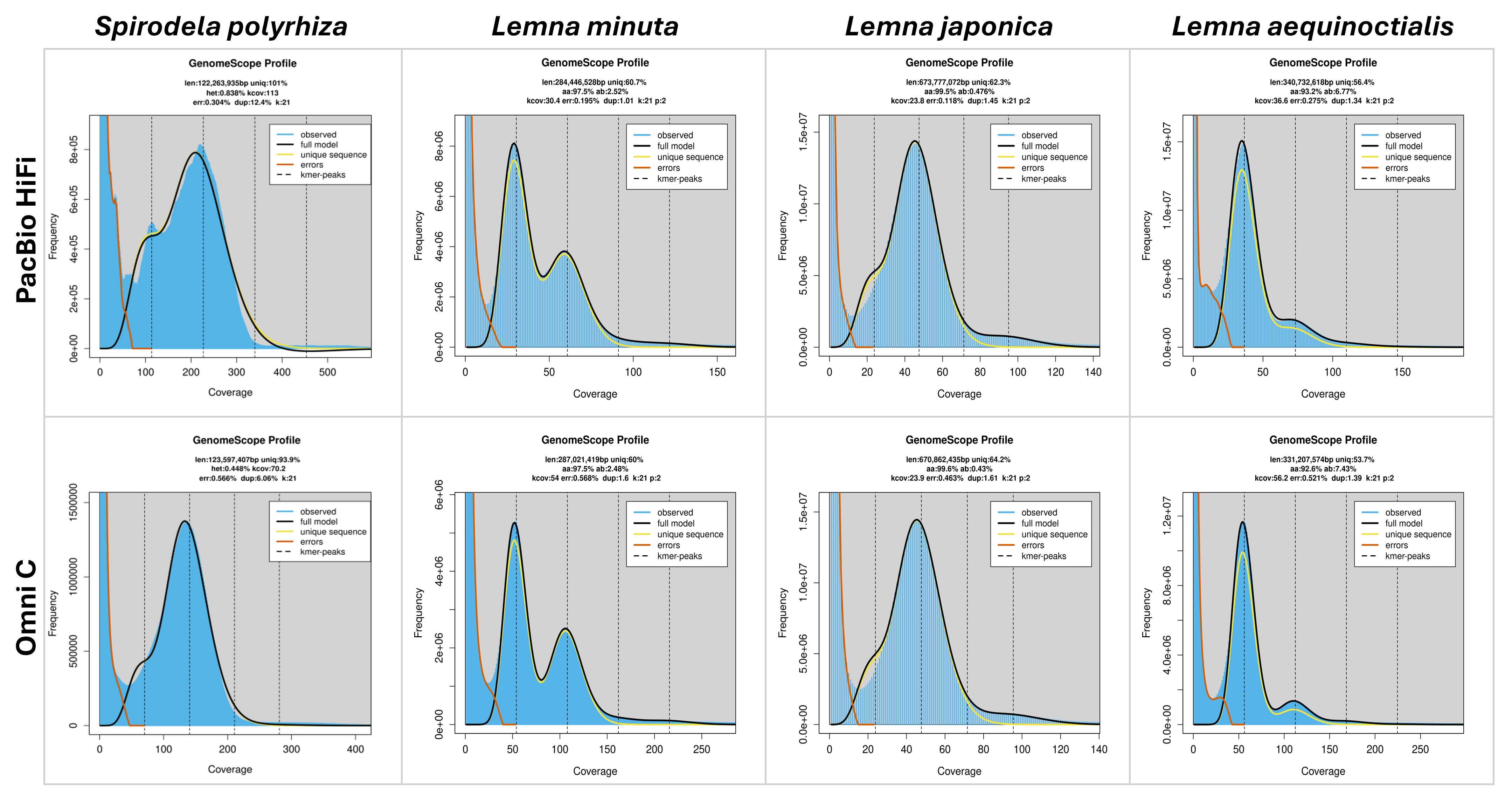

### Supplementary Figure 2

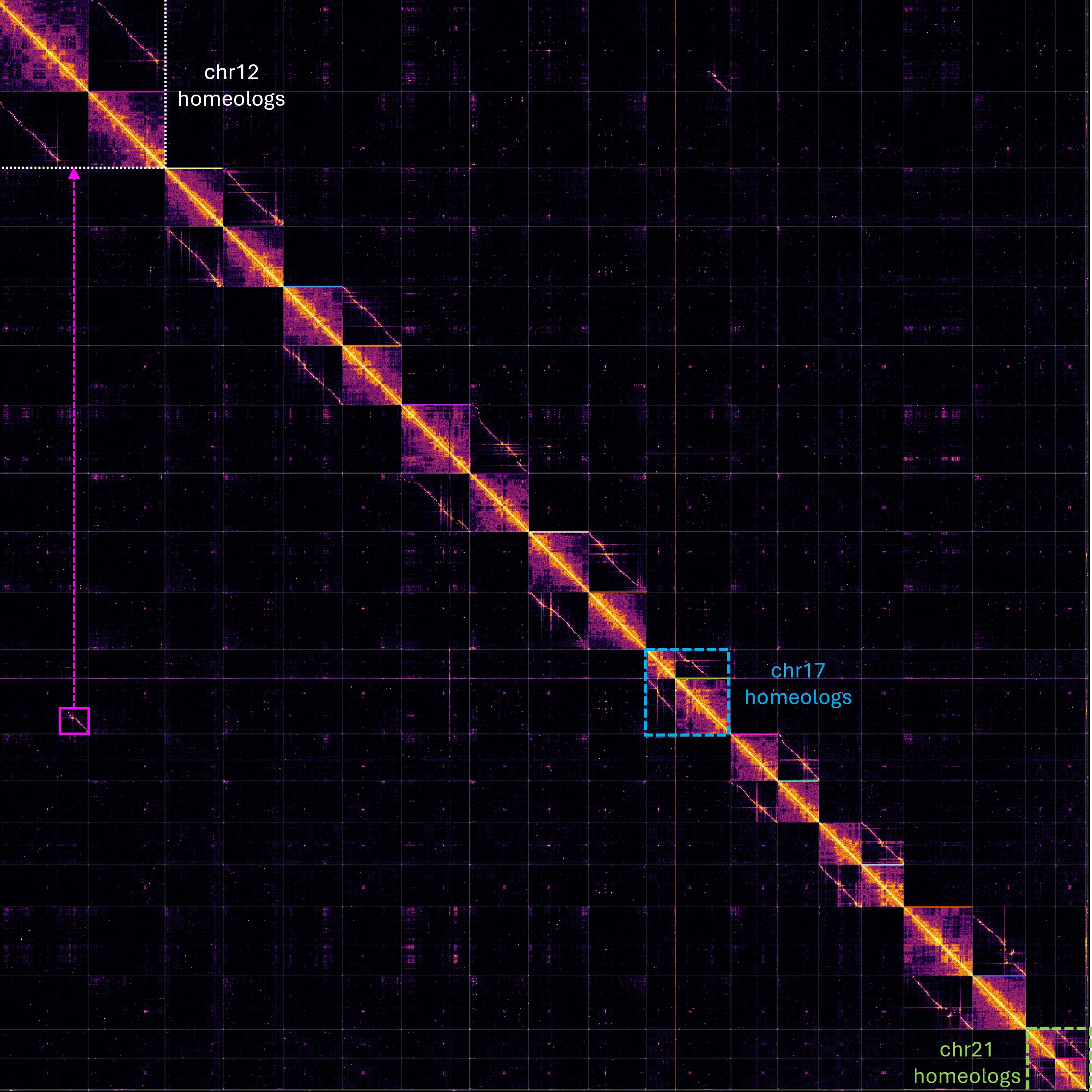

### Supplementary Figure 3

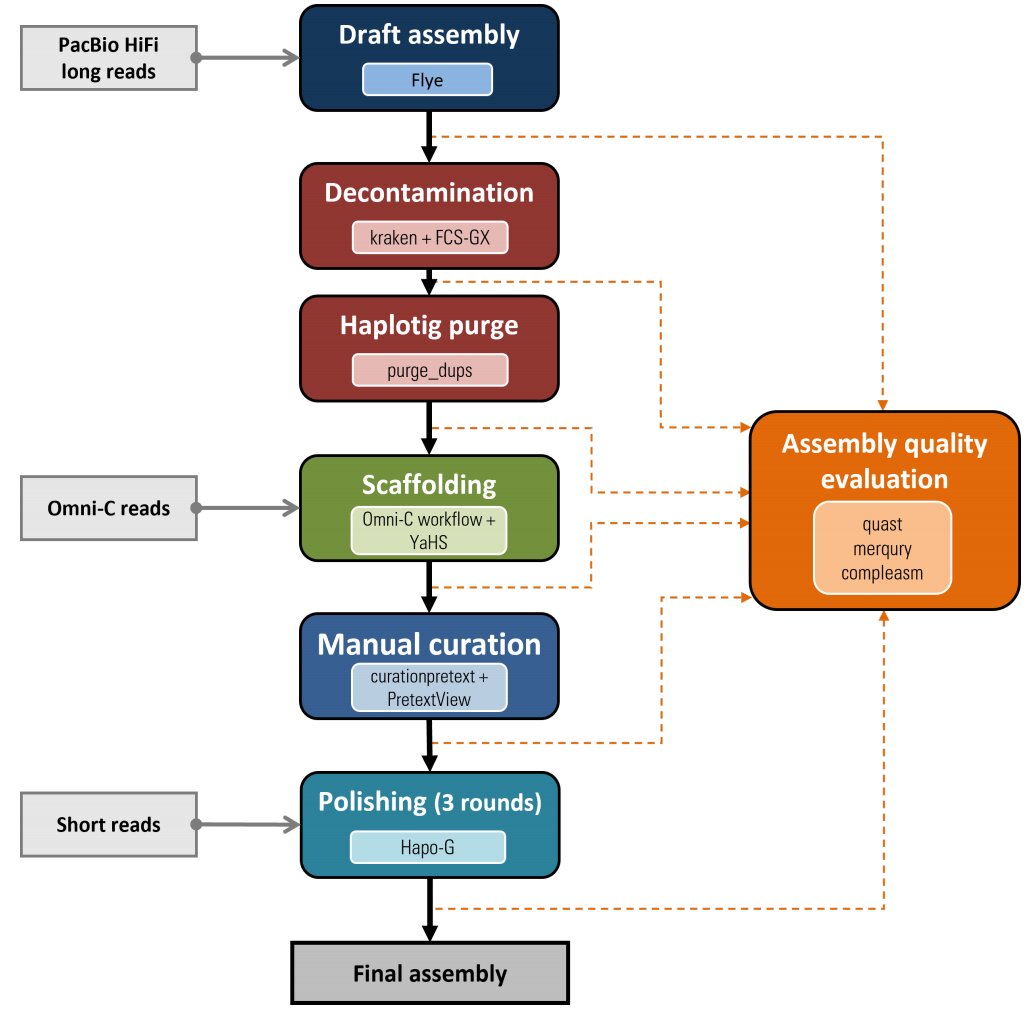

### Supplementary Figure 4

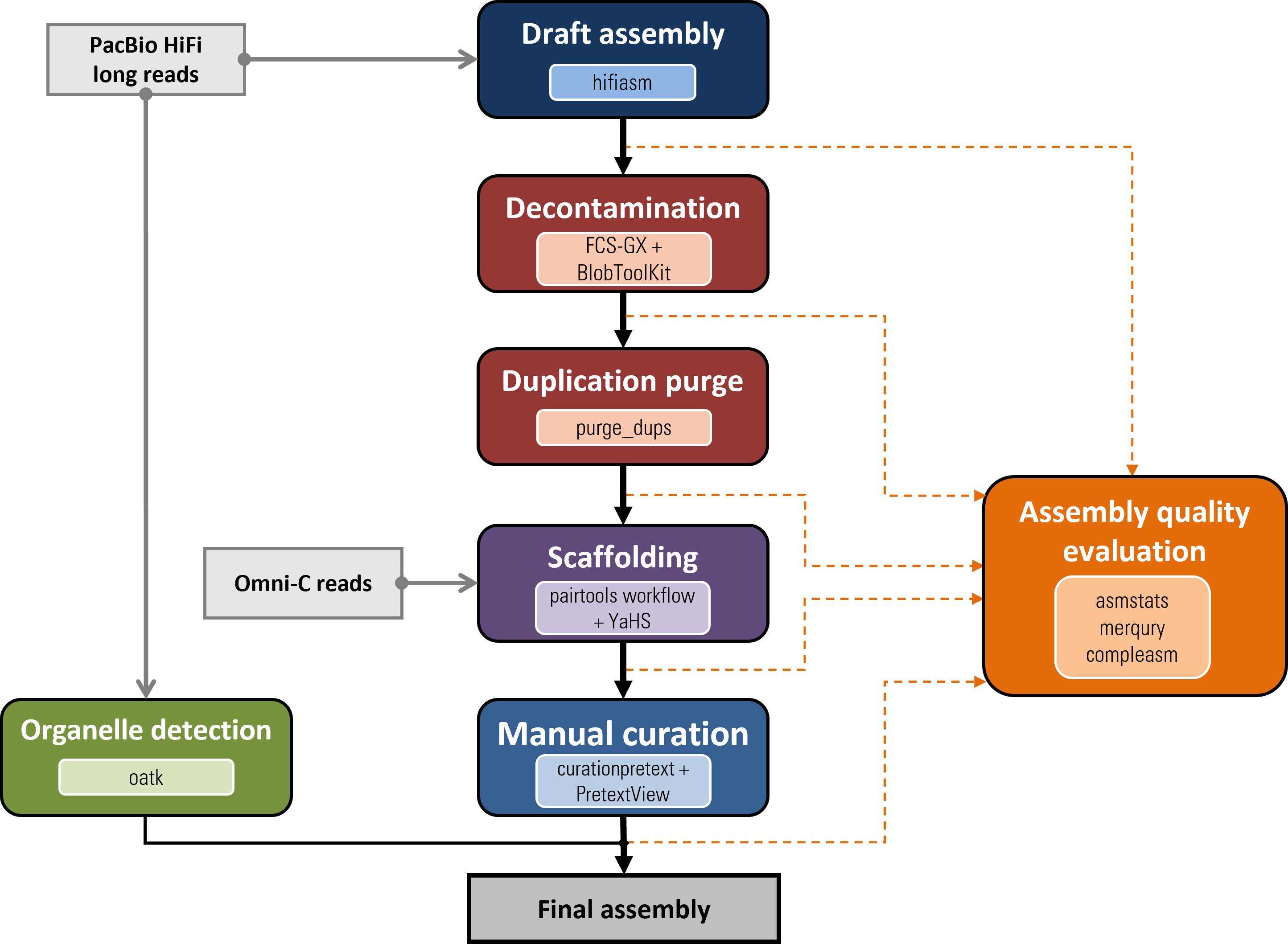

### Supplementary Figure 5

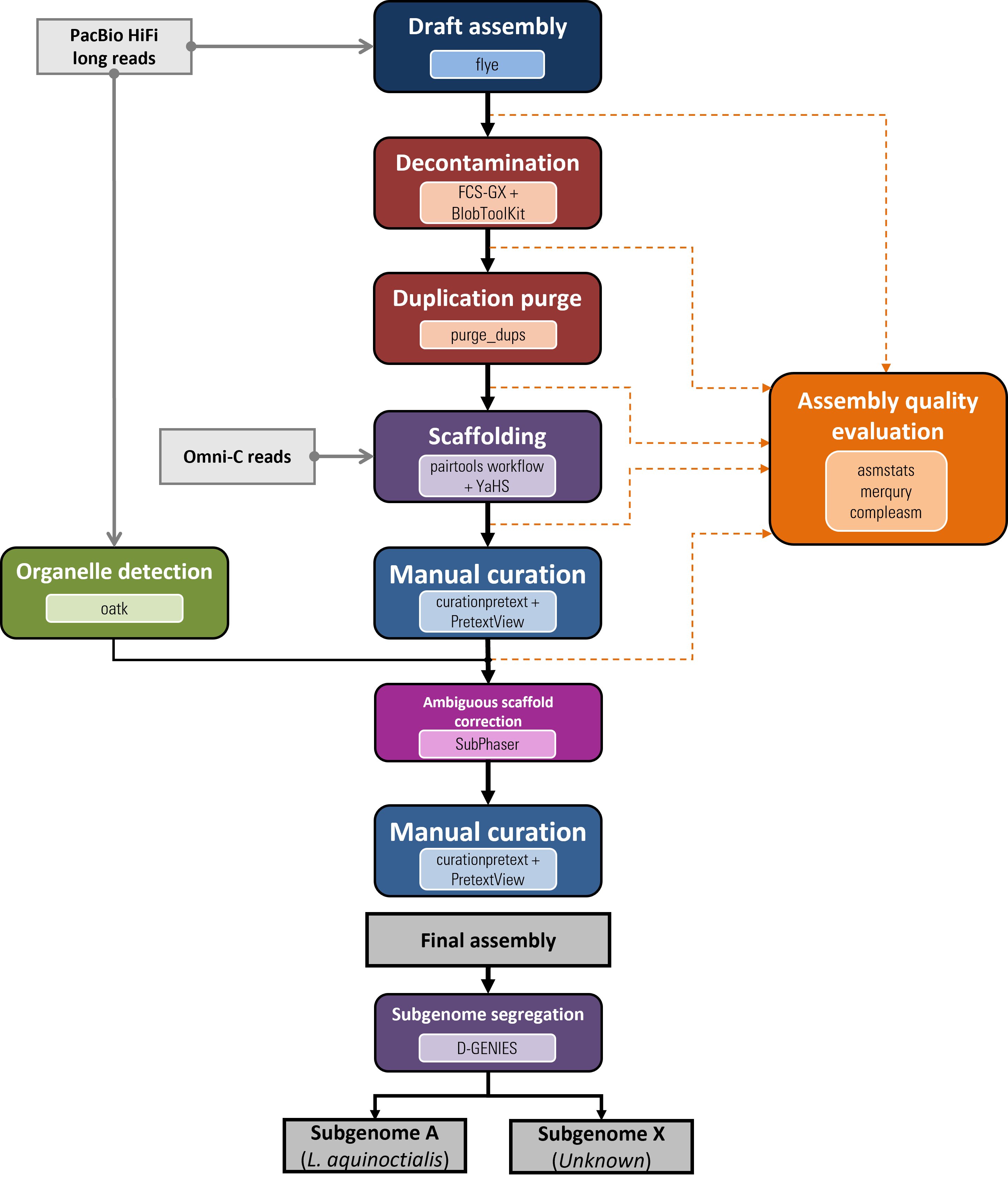

### Supplementary Figure 6

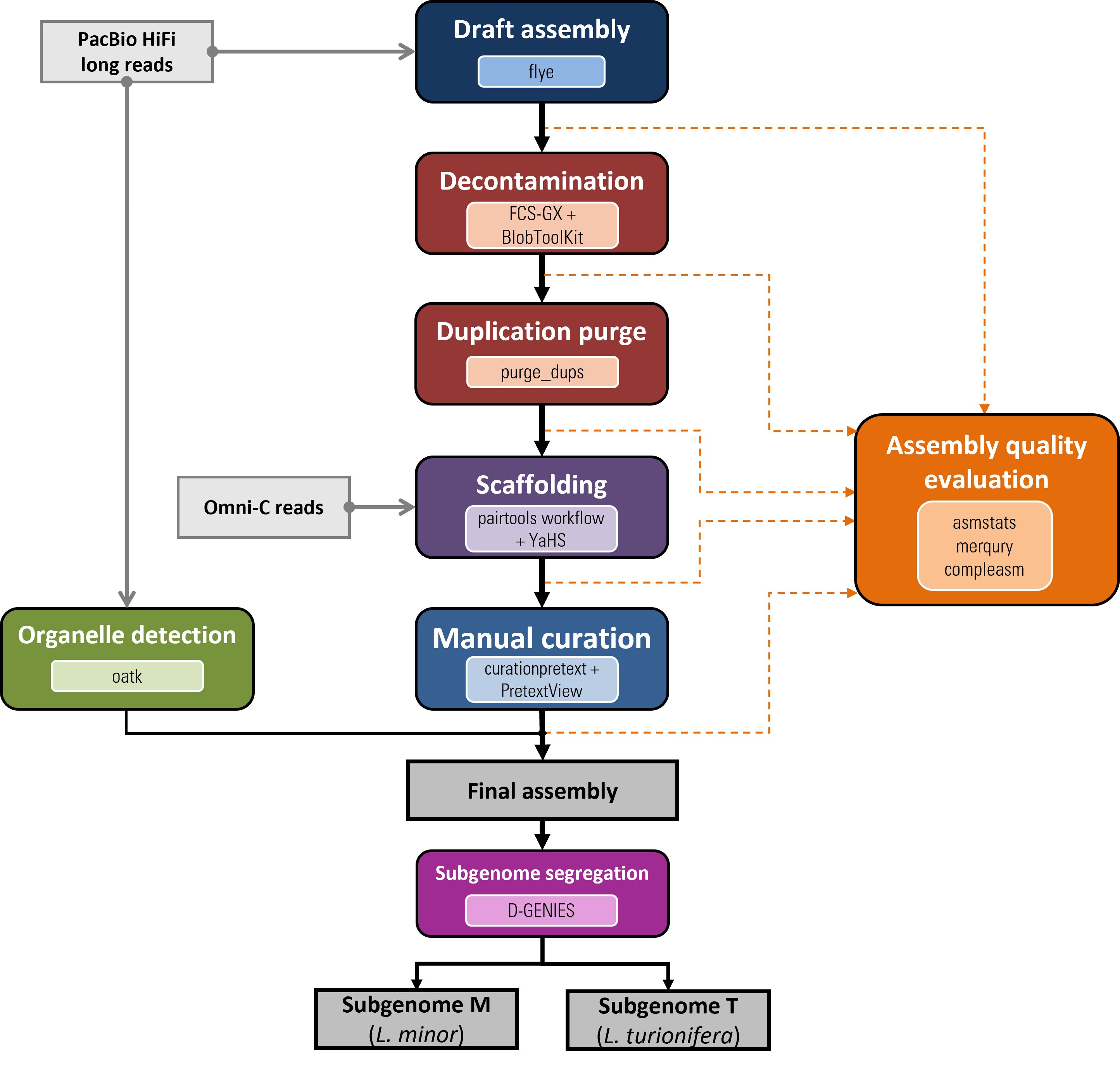

### Supplementary Figure 7

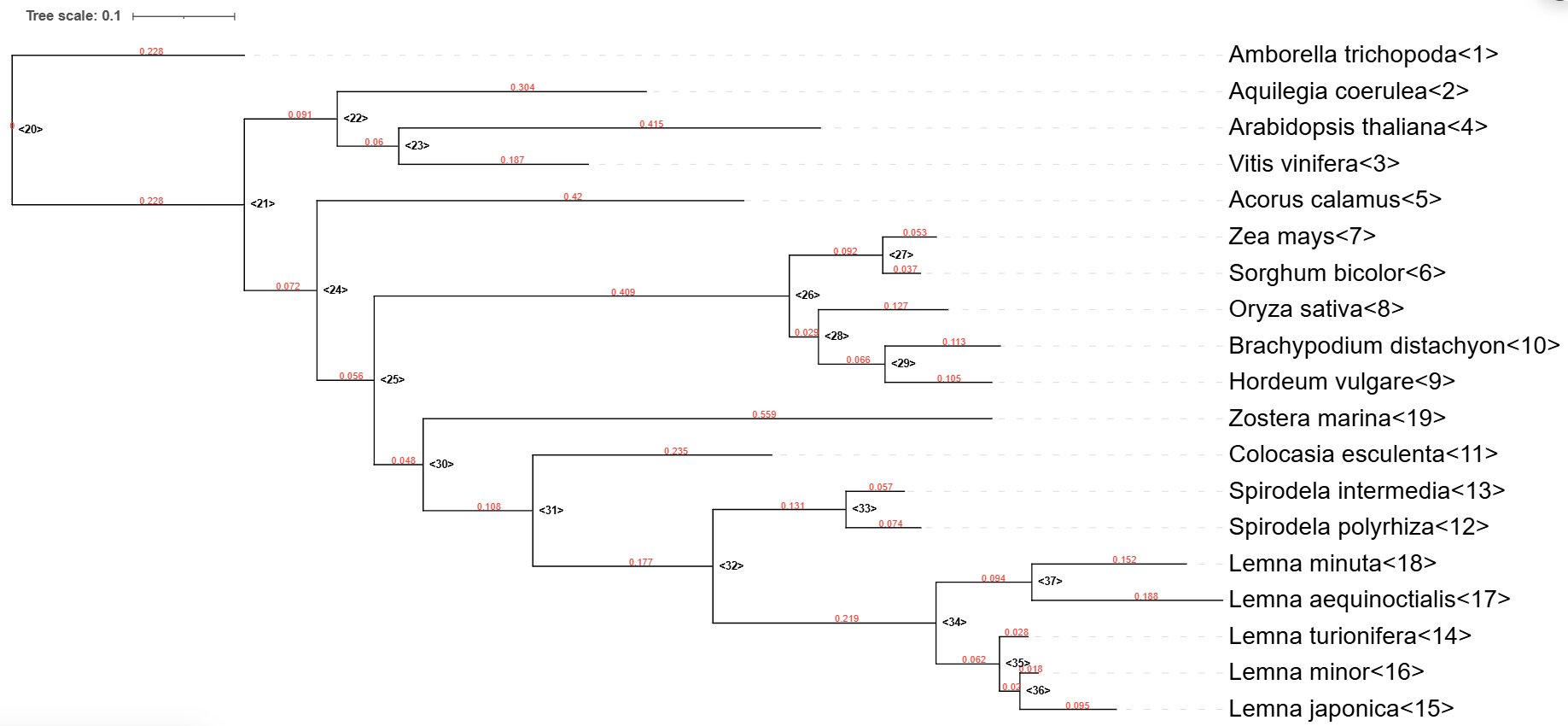

### Supplementary File 6

#### DEEPSPACE synteny map

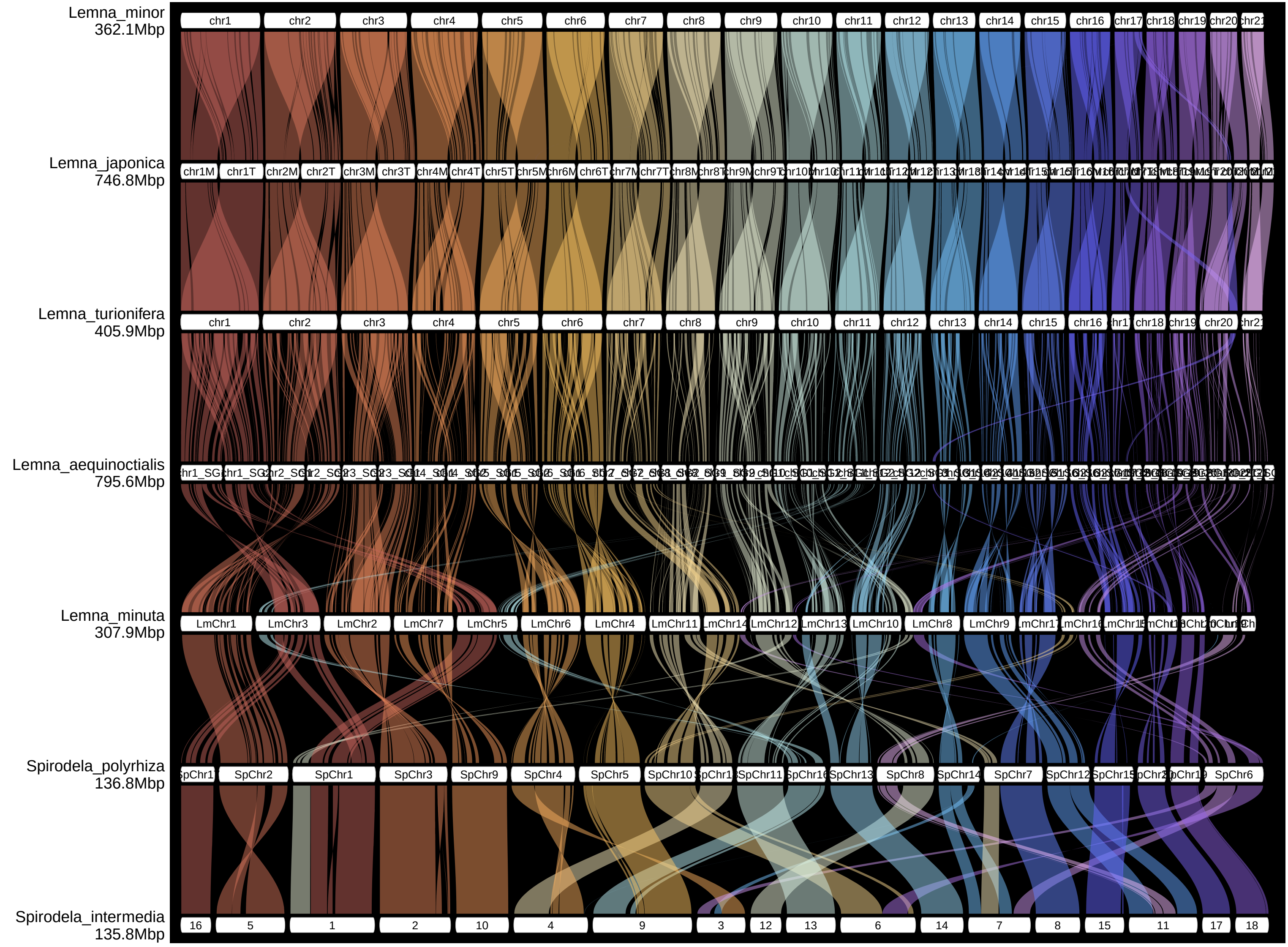

chromosomes scaled to the same x−range across genomes

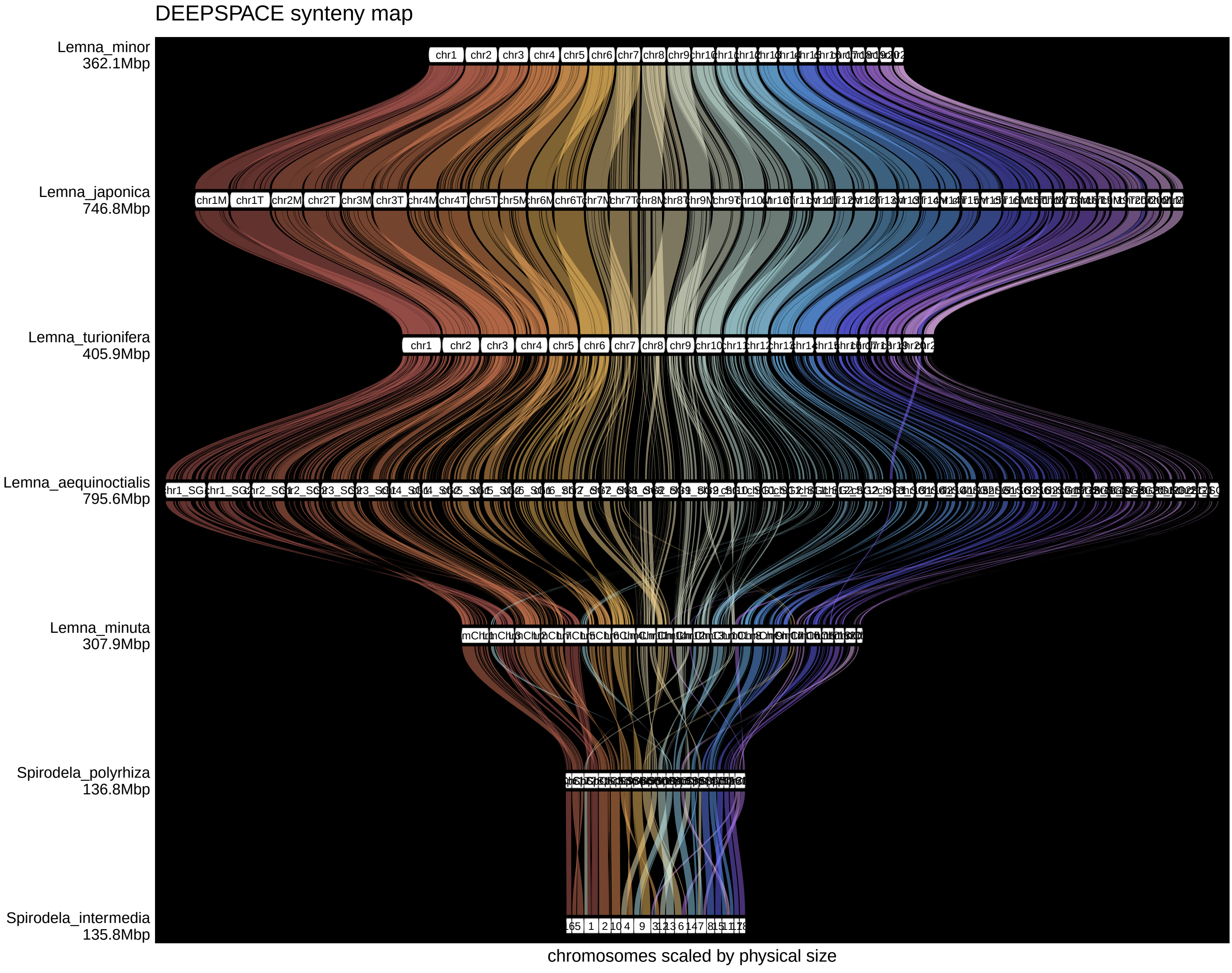

### Supplementary File 7

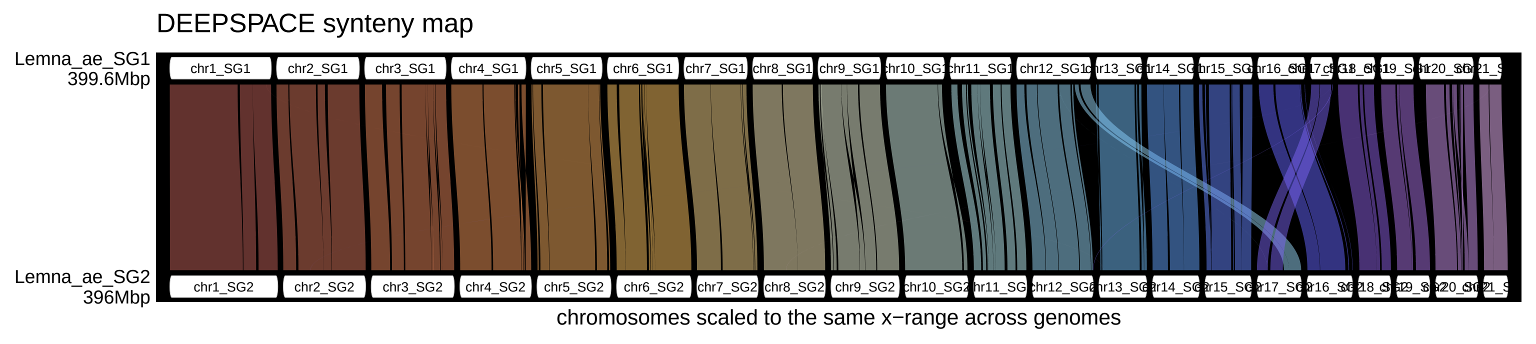

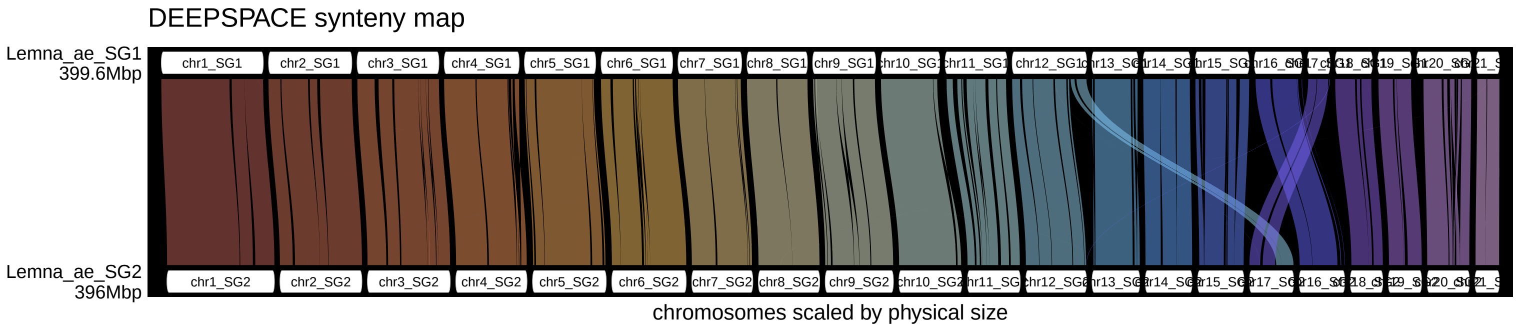
